# Single-cell RNA-sequencing implicates venous endothelial cells as a source of VEGF-A-mediated neo-angiogenesis in neuroinflammation

**DOI:** 10.1101/2022.11.15.516660

**Authors:** S. Shahriar, S. Biswas, K. Zhao, U. Akcan, M. C. Tuohy, M. D. Glendinning, A. Kurt, C. R. Wayne, G. Prochilo, M. Z. Price, R. A. Brekken, V. Menon, D. Agalliu

**Author notes:** Corresponding author, Mailing address: Columbia University Irving Medical Center, 650 West 168th Street, Black Building Room 310, New York, NY 10032. equal contribution first authors (S. Shahriar, S. Biswas). equal contribution senior authors (V. Menon, D. Agalliu).

## Abstract

Histopathological studies of multiple sclerosis (MS), a demyelinating disease of the central nervous system (CNS), and its animal model, experimental autoimmune encephalomyelitis (EAE), have found newly formed leaky vessels in demyelinated acute and chronic plaques, in addition to blood-brain barrier (BBB) damage in existing vessels, that exacerbate disease pathology by increasing infiltration of immune cells. Which vessel subtypes and signaling pathways generate these aberrant vessels is poorly understood. Using single-cell RNA-sequencing and *in vivo* validation, we find that transcriptome signatures of neo-angiogenesis arise in venous endothelial cells in both acute and chronic EAE, and correlate with upregulation in VEGF-A signaling. These neo-angiogenic markers are also increased in acute and chronic MS lesions. Treatment with a VEGF-A blocking antibody diminishes neo-angiogenic transcriptomic signatures and vascular proliferation *in vivo*, but does not restore BBB function or ameliorate significantly EAE pathology. Therefore, anti-angiogenic therapies in combination with immunomodulatory therapies may benefit MS progression.

## INTRODUCTION

Multiple sclerosis (MS) is a chronic, autoimmune, inflammatory demyelinating disease of the central nervous system (CNS), in which activated leukocytes invade the CNS to trigger neuroinflammation, demyelination, gliosis, and axonal damage and resection^1,2^. Inflammation of CNS endothelial cells (ECs) associated with focal breakdown of the blood-brain barrier (BBB) is prevalent in demyelinating MS plaques^3^. Abnormalities in BBB tight junctions precede the onset of clinical experimental autoimmune encephalomyelitis (EAE), the animal model for MS, as observed in histological^4–7^, MRI^8^, and *in vivo* two-photon imaging studies^9^. Endothelial tight junction abnormalities persist throughout the disease course and correlate with clinical severity in both MS^10,11^ and EAE^4,5^ due to their permissive role in leukocyte migration across the BBB^12–15^. Recent studies have shown that MS-associated single nucleotide polymorphisms are enriched in ECs compared to other CNS cells^16^, consistent with a critical role for the CNS endothelium in MS pathogenesis.

Angiogenesis is most active during CNS development in rodents and humans, and is absent in the adult CNS^17^. In addition to BBB damage in existing CNS vessels, neo-angiogenesis (the formation of abnormal new vessels) has been observed surrounding acute and chronic MS plaques^18,19^. Many blood vessels within MS/EAE lesions have an irregular morphology^20^, and express proliferation markers^21^. Vessel density is increased in demyelinating lesions during both peak and relapsing EAE, as well as in relapsing-remitting MS (RRMS) and secondary progressive MS (SPMS)^22–26^. A recent 3T SWI-FLAIR imaging study of the brain vasculature in RRMS patients using a 3T SWI-FLAIR also revealed several vascular abnormalities including small lesions at the vessel boundary, dilated vessels and developmental venous angiomas within MS lesions^27^. Most studies suggest that neo-angiogenesis worsens MS/EAE pathogenesis^18,19^ since its blockade improves clinical scores and attenuates demyelination and neuroinflammation in EAE^26,28^. However, the timing and relation of neo-angiogenesis to EAE/MS progression, and its molecular characterization, vascular origin, and underlying signaling pathways remain poorly understood.

Vascular endothelial growth factor A (VEGF-A) is a key mediator of angiogenesis in normal vascular development^29^, and in pathologies such as cancers^30^, ocular conditions^31^ and inflammatory diseases. VEGF-A expression is upregulated in acute and chronic MS plaques^20^ and EAE^26,28^. Serum VEGF-A levels are significantly higher in relapsing MS, compared to either remitting MS patients, or healthy cases, and correlate with lesion length on MRI examination^20,32^. Bevacizumab (Avastin®) is a recombinant, humanized monoclonal antibody that inhibits VEGF-A by blocking its binding to VEGF Receptor 2 (VEGFR2) and VEGFR1^33–35^. Administration of either bevacizumab, or the monoclonal antibody B20.4-1-1 that binds both murine and human VEGF-A^28^, in mice prior to and during EAE induction improves clinical disease and attenuates both demyelination and inflammation^26,28^. VEGF-A modulates also immune cell function and suppresses host immunity in the tumor microenvironment^36^. Critically, previous studies have failed to disentangle whether VEGF-A blockade ameliorates EAE clinical and histopathological outcomes by inhibiting angiogenesis, or by reducing peripheral immune cell activation and CNS infiltration.

In addition to VEGF-A signaling, the Wnt/β-catenin pathway is critical for both developmental angiogenesis and BBB formation and maintenance in the healthy CNS^37–41^. We have previously shown that the Wnt/μ-catenin pathway is reactivated in CNS ECs during EAE and MS. Its inhibition in CNS ECs in mice prior to EAE onset exacerbates disease outcomes, increases CD4^+^ T cell infiltration into the CNS, and exacerbates demyelination by increasing expression of Vascular cell adhesion molecule-1 (Vcam1) and Caveolin-1, two proteins critical for lymphocyte attachment and migration across the BBB^9,42^. Whether activation of the Wnt/μ-catenin signaling in EAE/MS promotes neo-angiogenesis together with VEGF-A remains unknown.

To address which vessel subtypes and signaling pathways drive neo-angiogenesis and characterize its molecular signatures in neuroinflammation, we performed single-cell RNA-sequencing (scRNA-seq), and subsequent *in vivo* validation in control, acute and chronic MOG_35-55_ EAE mice. We find that transcriptomic signatures of neo-angiogenesis are upregulated in venous ECs in both acute and chronic EAE, and correlate with upregulation in VEGF-A, but not Wnt/μ-catenin, signaling. These neo-angiogenic markers are also increased in acute and chronic MS lesions. Treatment with the VEGF-A blocking antibody r84^43,44^ diminishes neo-angiogenic transcriptomic signatures and reduces vascular proliferation *in vivo*, but fails to restore BBB function or significantly ameliorate EAE pathology. Our study provides a comprehensive molecular understanding of the origin and mechanism of neo-angiogenesis in EAE/MS, and suggests that anti-angiogenic therapies in combination with immunomodulatory therapies may benefit MS progression.

## RESULTS

### ECs isolated from spinal cords of control and EAE mice cluster based on A-V zonation and disease state

To determine which vessel subtypes and signaling pathways drive neo-angiogenesis in EAE, we induced MOG_35-55_ EAE in 12-week-old *TCF/LEF1::*H2B::eGFP (Wnt reporter) female mice^9,42^. We combined spinal cords from 2 mice for each sample, and collected the tissue at acute peak [score = 2-4; 16-17 days post-immunization (d.p.i.); n=8 mice, 4 samples] and early chronic (score ≤ 1; 28-29 d.p.i.; n=8 mice, 4 samples) disease for scRNA-seq (**Figure 1a**). Age-matched healthy (n=6 mice, 3 samples) and Complete Freund’s Adjuvant (CFA)-immunized mice (n=6 mice, 3 samples) served as controls. Acute and chronic EAE mice had a similar disease course with no difference in disease onset, or peak clinical score. The clinical score on the day of collection was higher in the acute compared to chronic cohort (CFA mice did not have any disease; **Extended Data Figure 1a-e**). Spinal cords were dissociated into single-cell suspensions and Wnt-positive (eGFP^+^) and Wnt-negative (eGFP^-^) CD31^+^ ECs were sorted and pooled for scRNA-seq (**Extended Data Figure 1f**). After quality filtering^45^ and *in silico* EC selection based on expression of *Pecam1*, *Cldn5*, and *Erg*^46^, we performed graph-based clustering to separate 45,309 ECs collected from from 14 samples [4 acute EAE, 4 chronic EAE, 3 healthy and 3 CFA controls (28 total mice, 2 mice/sample, **Extended Data Table 1-1**)] into distinct clusters based on their transcriptomic expression profiles.

**Figure 1.**
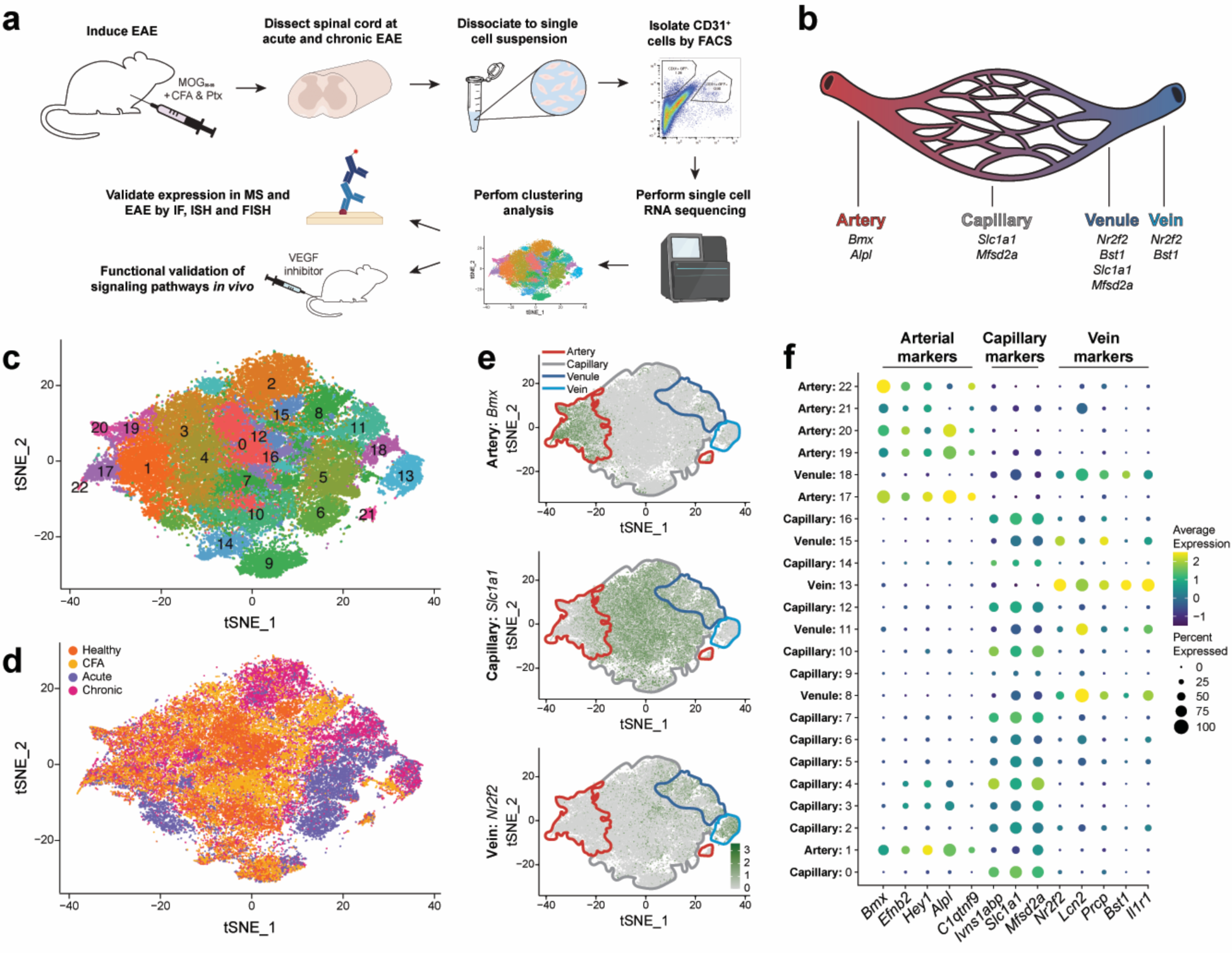
Single-cell RNA-sequencing and analysis of spinal cord endothelial cells from control and MOG_35-55_ EAE mice reveals clustering driven by zonation followed by disease/healthy state. **a)** Schematic diagram of the experimental workflow for the isolation and transcriptomic profiling of endothelial cells (ECs) from either healthy (orange), CFA (yellow), acute (blue) or chronic (magenta) MOG_35-55_ experimental autoimmune encephalomyelitis (EAE). We used two mice per sample to obtain sufficient ECs from spinal cords for scRNA-seq analysis. **b)** Architecture of the mammalian vascular system depicting arteries, capillaries, venules and veins and some of their specific molecular markers. **c)** t-SNE visualization of the 23 distinct EC clusters within the scRNA-seq dataset. **d)** t-SNE plot color-coded by sample condition. ECs from acute (n = 4 samples; blue) and chronic (n = 4 samples; magenta) EAE samples cluster separately from control ECs [healthy (n = 3 samples; orange) and CFA (n = 3 samples; yellow)], suggestive of transcriptional shifts in ECs from both acute and chronic EAE. **e)** Feature plots show the distribution of arterial (*Bmx*), capillary (*Slc1a1*), and venous (*Nr2f2*) marker expression within the t-SNE plot. Expression of arterial and venous markers peaks at the opposite ends of the t-SNE plot separated by the capillary marker expression in the middle indicative that zonation is an important axis of variation among the cells. **f)** Dot plot illustrating the expression of several arterial-, capillary-, and venous-specific markers within the 23 EC clusters in the scRNAseq dataset. Clusters are annotated based on marker gene expression.

Graph-based clustering at high resolution separated the ECs into 23 distinct clusters, which were visualized using a *t*-distributed stochastic neighbor embedding (t-SNE) representation (**Figure 1c-f)**. While these original 23 clusters are not necessarily functionally distinct, this “overclustering” approach was used to establish clear boundaries among cells belonging to each of the major vessel subtype. Overclustering was followed by remerging clusters defined by computational or biological measures; this two-step process can avoid suboptimal clustering boundaries that often arise when using lower resolution clustering^47^. After the merging step, we assigned a major subtype identity to each EC cluster based on expression of canonical markers for arteries, capillaries and veins^48^ (**Figure 1b**). Expression of arterial and venous markers peaked in clusters visualized at opposite ends of the t-SNE plot; putative capillary clusters organized themselves between these ends in t-SNE embedding (**Figure 1e, f**). Although non-local distances in t-SNE plots are highly nonlinear, the overall arrangement of ECs suggests that arteriovenous (AV) zonation was a major putative axis of variation in the EC data (**Figure 1b, e**). The disease state was the second putative axis of variation since ECs purified from acute and chronic EAE formed separate groups from those isolated from controls (**Figure 1c, d, Extended Data Figure 1i**). To validate the robustness of vascular subtype classifications for our single cell clusters, we compared p-value distributions of differentially expressed genes among subtypes from shuffled data where we randomly ordered the cell cluster assignments. The differential gene lists specific to vascular subtype cluster demonstrated substantially and significantly more hits than those obtained from random shuffling, an indication that despite the sparsity and dropouts inherent in of single-cell datasets, the signal retains biological relevance for vascular subtype identification (data not shown). Given that our samples were processed in separate batches (**Extended Data Table 1-1**), we conducted clustering with and without Harmony-based^49^ batch correction to assess the degree of batch effects (**Extended Data Figure 1g, h**). Comparing cluster membership before and after batch correction revealed insignificant batch impact on the membership of the four major vascular subtypes, signifying the robustness of vascular subtype clustering to batch effects. The minimal influence of batch on subtype clustering is additionally corroborated visually by the overlay of batch structure onto the individual vascular subtype-specific t-SNE plots (**Extended Data Figure 1i**). Finally, quantification of *eGFP* mRNA expression within the single-cell dataset revealed that Wnt/β-catenin signaling was highest in capillary ECs (cECs), moderate in arterial ECs (aECs) and venule ECs (vnECs), and low-to-absent in venous ECs (vECs) (**Extended Data Fig. 1j**).

### Transcriptomic signatures of neo-angiogenesis arise in venous ECs in both acute and chronic EAE

EC clusters belonging to each vascular subtype (6,856 aECs; 30,208 cECs; 6,596 vnECs, and 1,649 vECs) were subclustered separately and visualized again with t-SNE (**Figure 2a-h**). Arterial ECs from acute EAE formed a separate sub-cluster from chronic EAE and control samples which clustered together (**Figure 2a, b, i, j**), suggesting that transcriptomic changes induced by neuroinflammation are transient in aECs. In contrast, cECs, vnECs, and vECs isolated from control, acute, and chronic EAE formed distinct sub-clusters (**Figure 2c-j**), indicative of more persistent transcriptomic changes during neuroinflammation. The control (healthy and CFA)-enriched cluster in both cEC and vnEC populations contained a higher proportion of ECs originating from chronic versus acute EAE (**Figure 2i, j**). In contrast, the control-enriched cluster in vECs contained only a small percentage of ECs from chronic EAE (**Figure 2i, j**). Overall, the control samples (healthy and CFA) have similar transcriptional profiles, except in vnECs where they cluster separately due to increased expression of inflammatory gene signatures in CFA (**Figure 2e, f, i, j**; data not shown). Lastly, there were no observed differences in the proportions of aECs, cECs, vnECs, and vECs collected from both healthy and diseased samples (**Figure 2j**). These data confirm that ECs undergo transcriptional shifts in both acute and chronic EAE, consistent with prior reports using bulk and scRNA-seq^50,51^.

**Figure 2.**
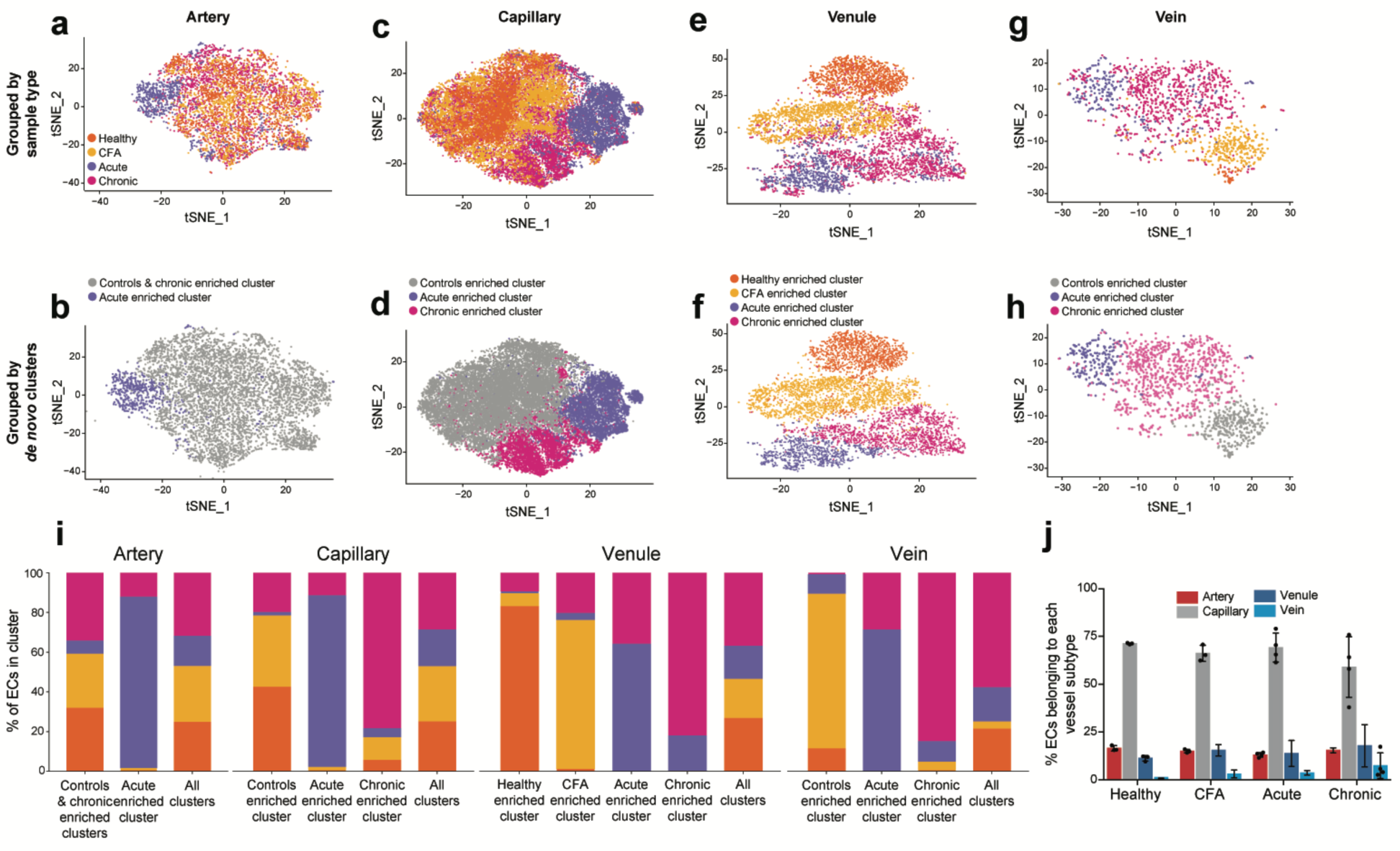
Endothelial subtype-specific t-SNE plots for spinal cords ECs isolated from control and EAE mice show separation of capillary and venous ECs in acute and chronic EAE from the healthy/CFA state. **a, c, e, g**) Individual t-SNE plots for arterial (aECs; a), capillary (cECs; c), venule (vnECs; e) and vein ECs (vECs; g) color-coded for sample type. **b, d, f, h**) Individual t-SNE plots for aECs (b), cECs (e), vnECs (h) and vECs (k), respectively, depict the clustering of *de novo* clusters for each vessel subtype. There is no clear separation in either the clustering or the t-SNE visualization between the two controls (healthy and CFA) and chronic EAE in aECs (colored in gray) and the two controls (healthy and CFA) in cECs and vECs (colored in gray). **i**) Stacked plots showing the percentage of ECs from each sample type (healthy, CFA, acute or chronic EAE) within the aECs (c), cECs (f), vnECs (i) and vECs (l) t-SNE plots, respectively. j) Dotted bar graphs show the proportion of arterial (aECs; red), capillary (cECs; gray), venule (vnECs; dark blue) and vein (vECs; light blue) ECs isolated from each sample type (healthy, CFA, acute and chronic EAE). Graphs represent mean ± SEM; *p < 0.05, **p < 0.01, **p < 0.001, ****p < 0.0001; one-way ANOVA with Tukey test.

To identify key transcriptomic signatures driving EC clustering, we performed differential gene expression (DE) analysis comparing the acute- and chronic-EAE to control-enriched EC clusters for each vessel subtype (**Extended Data Table 1-3 to 1-9**). This data was used to conduct gene set enrichment analysis (GSEA)^52^ using curated and comprehensive gene catalogs (**Extended Data Table 1-2**) for various cellular and molecular pathways and functions from the Molecular Signatures Database v6.1 (*MSigDB: software.broadinstitute.orggsea/msigdb/genesets.jsp*) and published sources^53,54^, and determine which processes were enriched for significantly up- and downregulated genes in diseased ECs from each vessel subtype (**Extended Data Table 1-3 to 1-9**).

Gene sets associated with inflammation, antigen processing and presentation, and chemokine/cytokine signaling were upregulated in all ECs from acute EAE (**Figure 3a**), and those associated with inflammation and antigen processing and presentation were also upregulated in cECs and vECs from chronic EAE (**Figure 3c**). In contrast, expression of the BBB gene set was significantly downregulated in diseased cECs, vnECs, and vECs in both acute and chronic EAE (**Figure 3b, d**), consistent with a dysfunctional BBB^50,51^. Surprisingly, expression of BBB genes was upregulated in acute aECs relative to controls (**Figure 3a**), which may explain the transient nature of neuroinflammatory transcriptomic changes in aECs (**Figure 2b**).

**Figure 3.**
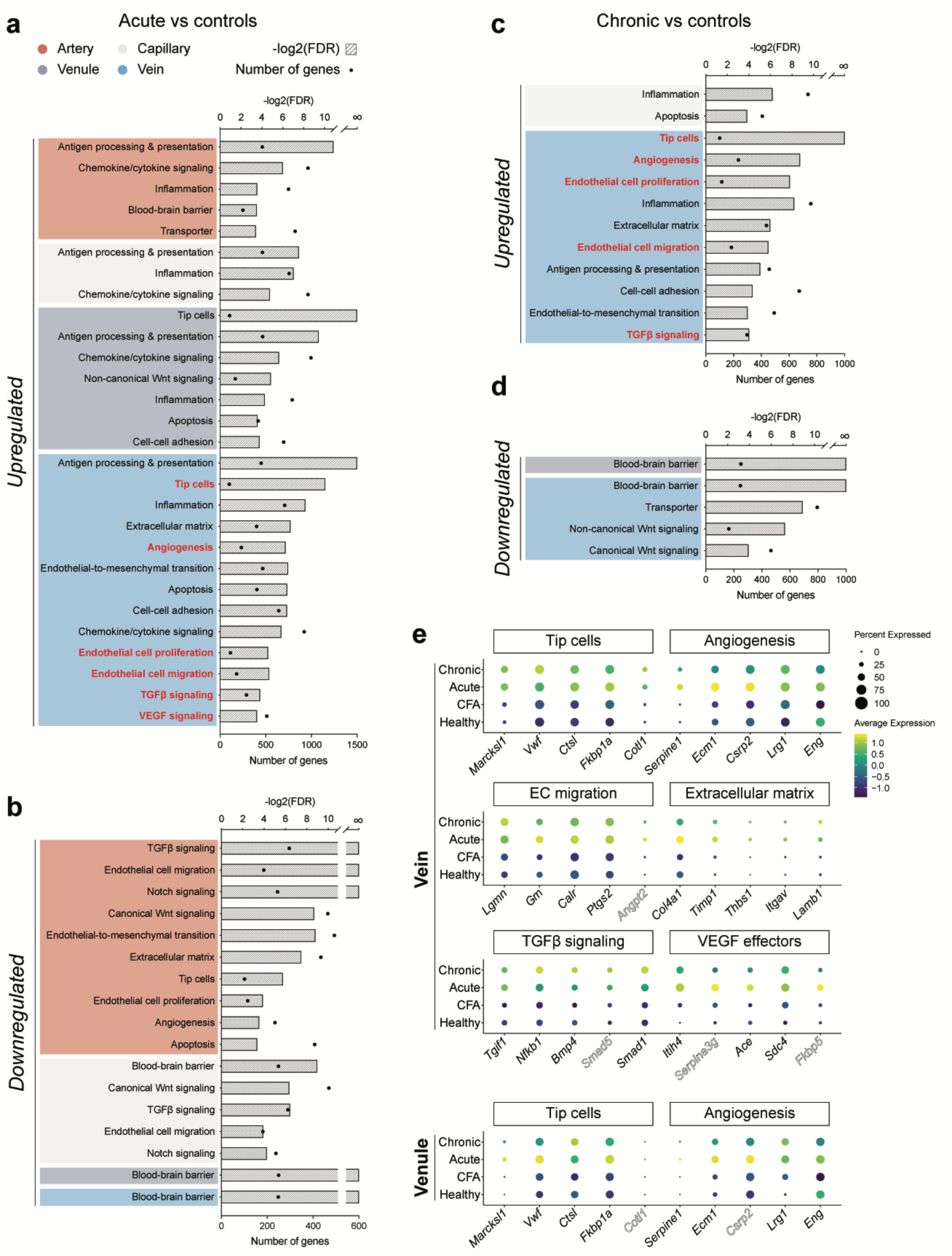
Gene set enrichment analysis (GSEA) reveals that transcriptome signatures of neo-angiogenesis are present in vein ECs from both acute and chronic EAE. **a, c**) Pathways/processes upregulated in diseased arterial (red), capillary (light gray), venule (dark blue), and venous (light blue) ECs in acute (a; 16 d.p.i) and chronic (c, 28 d.p.i) EAE versus controls. The gene ontology enrichment plots, derived using the GSEA algorithm, indicate that gene sets related to neo-angiogenesis (highlighted in red) are significantly upregulated in diseased vECs, and to a lesser extent, vnECs, relative to controls, in both acute and chronic EAE. **b, d**) Pathways/processes significantly downregulated in diseased arterial (red), capillary (gray), venule (venetian blue) and venous (pool blue) ECs in acute (b), and chronic (d), EAE versus controls. Expression of the blood-brain barrier gene set is significantly reduced in diseased ECs of all vessel subtypes, except arteries. **e**) Dot plots for expression of angiogenesis, tip cells, EC migration, extracellular matrix, VEGF and TGF-β pathway genes in vECs show increased average expression in both acute and chronic EAE. vnECs also upregulate expression of tip cell markers in acute and chronic EAE. Genes presented in solid typeface significantly upregulate during both acute and chronic EAE stages, whereas those in outlined font (Angpt2, Smad5, Serpina3g, Fkbp5, Cotl1, and Csrp2) show marked upregulation solely in the acute EAE phase.

Angiogenesis involves several steps ranging from tip and stalk EC specification, tip cell-mediated extracellular matrix (ECM) degradation, EC migration and proliferation^55^. Pathological tumor angiogenesis is also regulated, in part, by endothelial-to-mesenchymal transition (EndoMT), which induces delamination of resident ECs and subsequent acquisition of a mesenchymal phenotype with invasive/migratory properties^56,57^. DE analysis of diseased versus control vnECs and vECs revealed that the top-upregulated genes are enriched for angiogenic and tip cell markers (features of neo-angiogenesis) in both acute and chronic EAE (**Figure 3e**; **Extended Data Figure 2**). Several angiogenesis (“tip cells”, “angiogenesis”, “EC proliferation” and “EC migration”) and angiogenesis-related (“ECM” ^58^, “EndoMT” ^59^) gene signatures were significantly upregulated in diseased veins (acute and chronic) together with two pro-angiogenic pathways (“VEGF” and “TGF-β”)^60–62^ (**Figure 3a, c**). Acute vnECs also showed upregulation of gene signatures for tip cells in acute EAE (**Figure 3a**). In contrast, arterial or capillary ECs in acute EAE were not enriched for transcriptional gene signatures of neo-angiogenesis compared to controls. In fact, expression of angiogenesis-related gene sets were downregulated in diseased aECs relative to controls, suggesting a vascular repair process and inverse correlation between BBB and angiogenesis-related genes (**Figure 3b**). Although several angiogenesis pathways were upregulated in EAE vECs, the Wnt/β-catenin signaling was almost absent in healthy vECs and downregulated in chronic EAE vECs (**Figure 3d; Extended Data Figure 1j**). In summary, our scRNAseq data reveal that transcriptomic signatures for neo-angiogenesis are upregulated in vECs, and to a lesser extent in vnECs, in acute and chronic EAE. While Wnt/β-catenin signaling is essential for developmental CNS angiogenesis^17^, it is likely not associated with pathological neo-angiogenesis in EAE.

### Congruency analysis reveals a unique transcriptome signature of neuroinflammation-related angiogenesis in EAE

We next asked to what extent neuroinflammation-related neo-angiogenesis shares similar transcriptomic signatures with either CNS developmental or tumor angiogenesis. We analyzed publicly available datasets of murine lung tumor Ecs (TECs)^63^ and developmental CNS Ecs [DECs; embryonic day 14.5 (E14.5)]^64^ to identify angiogenic genes that were significantly upregulated in expression during tumor and CNS developmental angiogenesis, respectively. Comparing those gene sets to the angiogenic gene signature of acute EAE vECs revealed that most angiogenic genes in neuroinflammation were also upregulated in developmental (∼68%) and tumor (∼85%) angiogenesis (**Extended Data Figure 3a-b**). There was a greater number of acute EAE vEC genes shared with TECs than with DECs, suggesting that neuroinflammation angiogenesis is more similar at the molecular level with tumor angiogenesis. We also analyzed the congruency in angiogenic gene expression between DECs and TECs (**Extended Data Figure 3c**). While pathogenic and developmental angiogenic gene signatures had many common genes (105), they also had a significant number of unique genes (DECs had 102 unique genes and TECs had 79), suggesting that the two processes are substantially different. Therefore, neuroinflammation and tumor angiogenesis have a higher congruency at the gene signature level, compared to neurodevelopmental angiogenesis.

To confirm these computational analyses *in vivo*, we analyzed expression of Apelin (*Apln*), one of the non-congruent angiogenic markers identified through the computational analysis between DECs and EAE vECs, in developing (P3), healthy adult, and acute EAE spinal cord tissues via mRNA *in situ* hybridization (**Extended Data Figure 3d-i**). *Apln* is a positive modulator of angiogenesis and is enriched in proliferating Ecs and tip cells in the P7 brain^65^. *Apln*^-/-^ mouse retinas show delayed EC proliferation, vascular outgrowth and branching^66^. Consistent with prior studies^65^, *Apln* mRNA was highly expressed by many developing blood vessels in the P3 spinal cord; however, its expression was abolished in the healthy adult CNS, except in the vasculature of the choroid plexus and meninges, and in some neurons (**Extended Data Figure 3d-i**; data not shown). *Apln* expression was not induced in inflamed blood vessels near white matter lesions of acute EAE spinal cords (**Extended Data Figure 3d-i**), validating that neurodevelopmental and neuroinflammatory angiogenesis are distinct at the transcriptome level.

### Venous ECs undergo proliferation and express angiogenesis markers in acute and chronic EAE

To validate the neo-angiogenic transcriptome signature observed in vECs in EAE (**Figure 3a, c**), we examined the expression of the proliferation marker Ki67 in arteries, capillaries, and veins in longitudinal (sagittal) spinal cord sections from control (CFA), acute and chronic EAE mice (**Figure 4a**). We used EphB4 as a marker for venous and venular ECs^67–69^, and Caveolin-1 to label all ECs. Consistent with scRNA-seq data for the *Mki67* transcript (**Figure 4b**), vECs in both acute and chronic EAE spinal cords showed significantly higher proliferation (Ki67^+^ ECs) compared to CFA controls (**Figure 4c, d**). The vascular coverage of EphB4^+^ vECs and venule diameter were increased significantly in acute and chronic EAE compared to the CFA spinal cords (**Figure 4c-f**). These data indicate that increased vEC proliferation in EAE, rather than sprouting angiogenesis, leads to venous expansion (diameter and coverage). Interestingly, EC proliferation (K67^+^ ECs) was absent in αSMA^+^ CD31^+^ aEC and Mfsd2a^+^ cEC in both control and EAE spinal cords (**Extended Data Figure 4a, b**). Therefore, EC proliferation is restricted to vECs in EAE.

**Figure 4.**
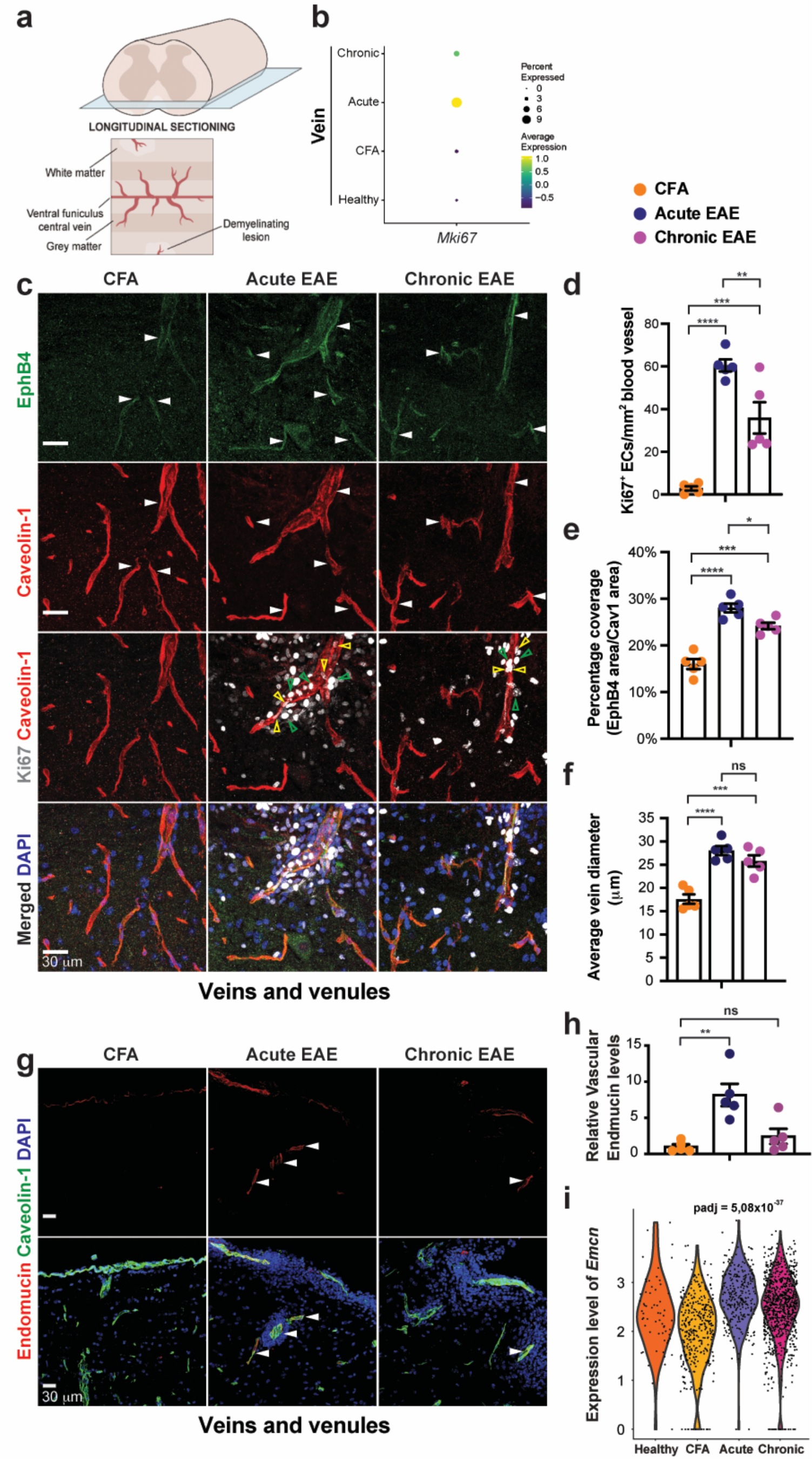
Increased vein EC proliferation in EAE leads to enlarged veins and higher venous coverage. **a)** Schematic diagram of the spinal cord longitudinal (sagittal) sections used throughout the study to measure *in vivo* protein/mRNA expression in control and EAE. **b)** Dot plot generated from the scRNA-seq dataset shows that *Mki67* mRNA expression is upregulated in acute, and to a lesser extent chronic, vECs. **c)** Sagittal sections of control (CFA), acute (16 d.p.i) and chronic (28 d.p.i) EAE spinal cords were immunostained with antibodies against EphB4 (venous ECs) and Ki67 (proliferation) along with Caveolin-1 (EC marker) and DAPI. White arrowheads point at EphB4^+^/Caveolin-1^+^ (double positive) vessels. Empty yellow arrowheads point at Ki67^+^ ECs. Empty green arrowheads point at Ki67^+^ immune cells. **d, e, f)** Dotted bar graphs showing quantification of number of Ki67^+^ ECs in EphB4^+^ vessels (d), EphB4^+^ vascular area (e) and vein diameter (f) reveal significantly higher EC proliferation, venous coverage and vein diameter in acute and chronic EAE spinal cords compared to the control (n = 5). **g)** Sagittal sections of control (CFA), acute EAE and chronic EAE spinal cords were immunostained for Endomucin (marks venous ECs), combined with immunostaining for Caveolin-1 (EC marker) and DAPI. White arrowheads point at expression of Endomucin in blood vessels. **h)** Dotted bar graphs showing relative expression level of Endomucin reveal significantly higher expression level in acute EAE spinal cord ECs compared to the control (n = 5). **i)** Violin plots for *Emcn* mRNA expression in control (healthy and CFA), acute and chronic EAE vECs generated from the scRNA-seq dataset. Graphs represent mean ± SEM from n = 5 mice/group; *p < 0.05, **p < 0.01, **p < 0.001, ****p < 0.0001; one-way ANOVA with Tukey test. Scale bars = 30 µm.

Endomucin (Emcn) is another integral membrane glycoprotein expressed in cECs and vECs^70^ implicated in modulation of VEGF-A-dependent angiogenesis in the retina^71^. Similar to EphB4, Emcn protein was increased in acute EAE spinal cord, relative to control and chronic EAE **(Figure 4g, h)**. These data are consistent with the analysis of differentially expressed genes by scRNAseq showing upregulation in *Emcn* expression in EAE vECs **(Figure 4i)**, but downregulated in diseased cECs. In contrast to ECs, there were no Ki67^+^ pericytes around blood vessels either in control or EAE spinal cords **(Extended Data Figure 4c)**.

Together, these data support our hypothesis that angiogenesis is unique to vECs in neuroinflammation. The lower vein density observed in chronic compared to acute EAE could be explained, in part, by a sequential specification of a venous (acute) followed by capillary and/or arterial fate (chronic) during disease, similar to postnatal developmental angiogenesis in the brain, or pathological angiogenesis in the retina^72–74^.

Differential gene expression analysis of diseased versus control vECs showed that the top upregulated transcripts in both acute and chronic EAE were angiogenesis and tip cell markers (**Figure 3e**). To validate these scRNA-seq findings *in vivo*, we analyzed expression of four key angiogenic/tip cell markers (*Egfl7*, Mcam, *Ecm1* and *Serpine1*) in spinal cords by fluorescent *in situ* hybridization (FISH) and immunofluorescence staining (IF). Epidermal growth factor-like 7 (Egfl7) is an ECM-associated protein that promotes angiogenesis in various organs including postnatal brain and retina^75–78^. Differential expression analysis of *Egfl7* expression by scRNA-seq did not show a significant upregulation in diseased relative to control vECs (**Figure 5a**). *Egfl7* mRNA levels were low-to-absent in CFA, but upregulated in inflamed blood vessels near white matter lesions in acute EAE using FISH combined with IF for Caveolin-1 (EC marker; **Figure 5e, i**). Melanoma cell adhesion molecule (Mcam), can function as a receptor for Wnt5a, Netrin1, FGF4, VEGF-C, and Wnt1^79^, or co-receptor for VEGF-A (together with VEGFR2)^80^ to promote angiogenesis. Mcam expression is enriched in proliferating ECs and tip cells of the brain and retina^65^. DE analysis by scRNA-seq showed significant upregulation of *Mcam* mRNA in both acute and chronic EAE vECs (**Figure 5b**). Mcam protein expression was also significantly upregulated in inflamed veins in acute EAE relative to CFA *in vivo* (**Figure 5f-j**). Extracellular matrix protein 1 (Ecm1) is a secreted glycoprotein that exerts its angiogenic function by potentiating epidermal growth factor receptor (EGFR) signaling^81^, which regulates synthesis and secretion of other angiogenic factors, including VEGF-A^82^. Differential expression analysis revealed that *Ecm1* mRNA was one of the most significantly upregulated genes in acute, and to a lesser extent, in chronic EAE vECs (**Figure 5c**). Assessment of *Ecm1* mRNA expression in control and EAE spinal cord tissue confirmed significant upregulation in acute EAE veins (**Figure 5g-k**). Finally, the serine protease inhibitor Serpine1 is a specific inhibitor of the plasminogen activation system^83^ and plays a crucial role in tumor progression and angiogenesis^84^. Expression of *Serpine1* was also significantly upregulated in acute, and to a lesser extent, chronic EAE vEC (**Figure 5d**). Assessment of *Serpine1* RNA expression *in vivo* revealed a trend toward increased expression in inflamed white matter blood vessels in acute EAE relative to control (p = 0.1261; **Figure 5h-l**). In summary, the *in vivo* validation of four angiogenic/tip cell marker expression identified from scRNA-seq confirms that neo-angiogenic genes are highly upregulated in inflamed veins in white matter lesions during peak (acute) EAE, but downregulated in chronic disease.

**Figure 5.**
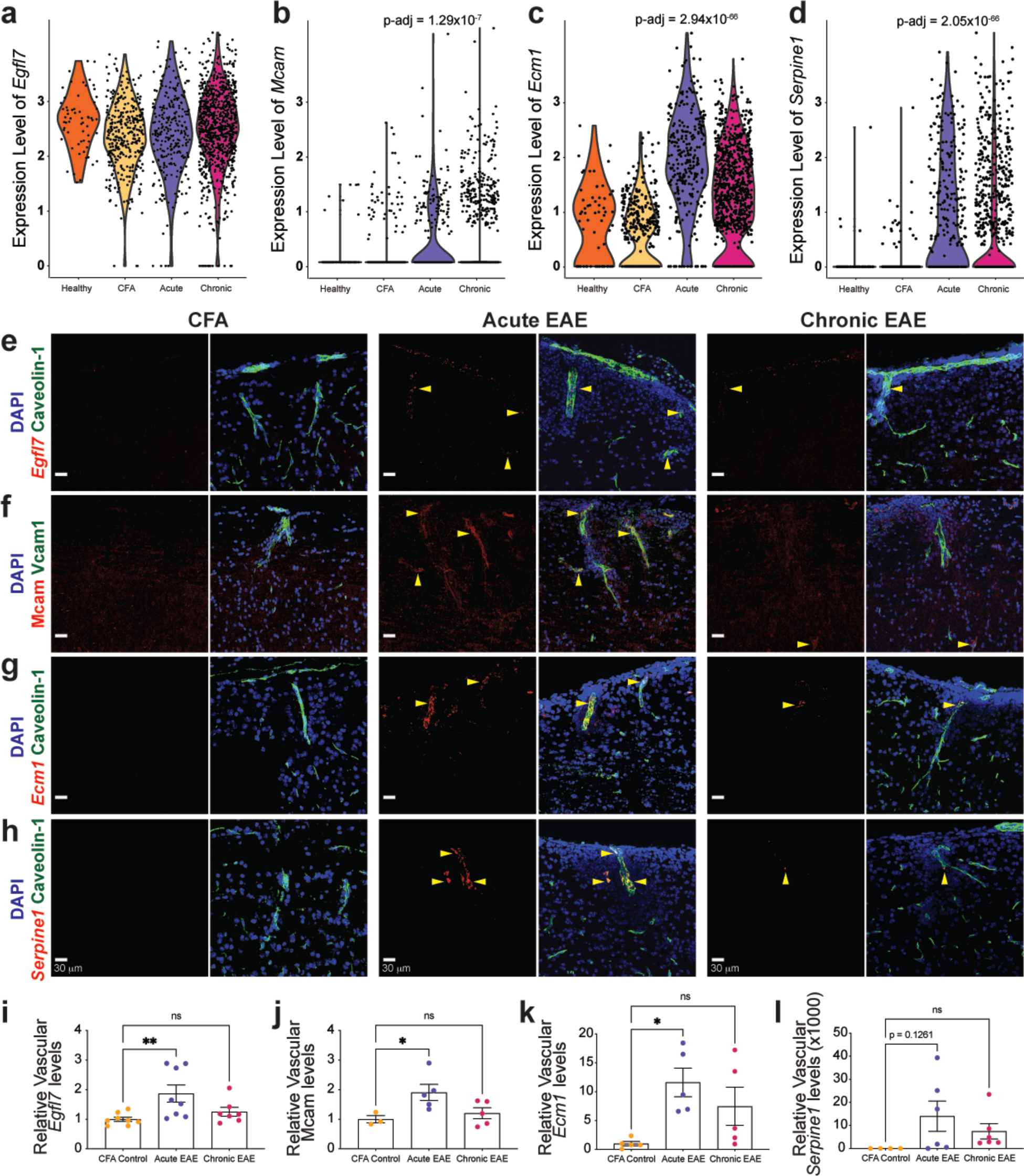
Expression of several tip cell/angiogenesis markers are upregulated by ECs in acute EAE. a-d) Violin plots for *Egfl7*, *Mcam*, *Ecm1* and *Serpine1* mRNA expression in control (healthy and CFA), acute and chronic EAE vECs generated from the scRNAseq dataset. e, f, g, h) Sagittal sections of control (CFA), acute and chronic EAE spinal cords were probed with fluorescence *in situ* hybridization (FISH) for *Egfl7* (e), *Ecm1* (g) and *Serpine1* (h) mRNAs or immunostained for Mcam (f), combined with immunostaining for Caveolin-1 (EC marker) or Vcam1 (venous marker) and DAPI. Yellow arrowheads point at the expression of corresponding markers in blood vessels. i, j, k, l) Dotted bar graphs showing relative expression levels of *Egfl7* (i), Mcam (j), *Ecm1* (k) and *Serpine1* (l) reveal significantly higher expression levels in acute EAE spinal cord ECs compared to the control. Data represent mean ± SEM; *p < 0.05, **p < 0.01, ****p < 0.0001; one-way ANOVA with Bonferroni’s multiple comparisons test. Scale bars = 30 µm.

### Expression of several angiogenic markers is upregulated in human MS tissue

Previous studies have shown higher vessel density and vessel abnormalities around acute and chronic MS plaques^27,85^, with irregular EC morphology^20^ and proliferation^21^. However, the molecular mechanisms of neo-angiogenesis in human MS are unknown. We examined expression of several angiogenesis markers, identified from scRNA-seq of Ecs in EAE, in acute or chronic demyelinating plaques within white matter (WM) lesions from16 MS patients, and compared them to either normal appearing white matter (NAMW) within the same tissue, or NAWM from brain samples of 16 non-neurological cases (**Extended Data Table 4**) with immunohistochemistry (12 cases per group), or *in situ* hybridization (4 cases per group). The CD31^+^ vascular coverage (vessel density) was slightly, albeit not significantly, higher inside the WM lesions of MS patients compared to NAWM in non-neurological controls (**Figure 6a-a”, d**). However, expression of two angiogenesis markers EGFL7 and MCAM proteins, was increased in blood vessels within WM lesions of MS patients compared to NAWM in non-neurological controls (**Figure 6b-c”, e-f**). There was a 4-fold increase in vascular EGFL7 expression and a 2-fold increase in MCAM^+^ vessels in WM lesions from chronic RMMS patients compared to the NAWM of non-neurological controls (**Figure 6e, f**). Expression of mRNAs for *SERPINE1* and *ECM1*, showed a similar pattern to EGFL7 protein (**Extended Data Fig. 5b-c”**). Thus, several molecular markers of pathological angiogenesis, identified from scRNA-seq of Ecs in EAE, are upregulated in demyelinating MS plaques in RRMS cases.

**Figure 6.**
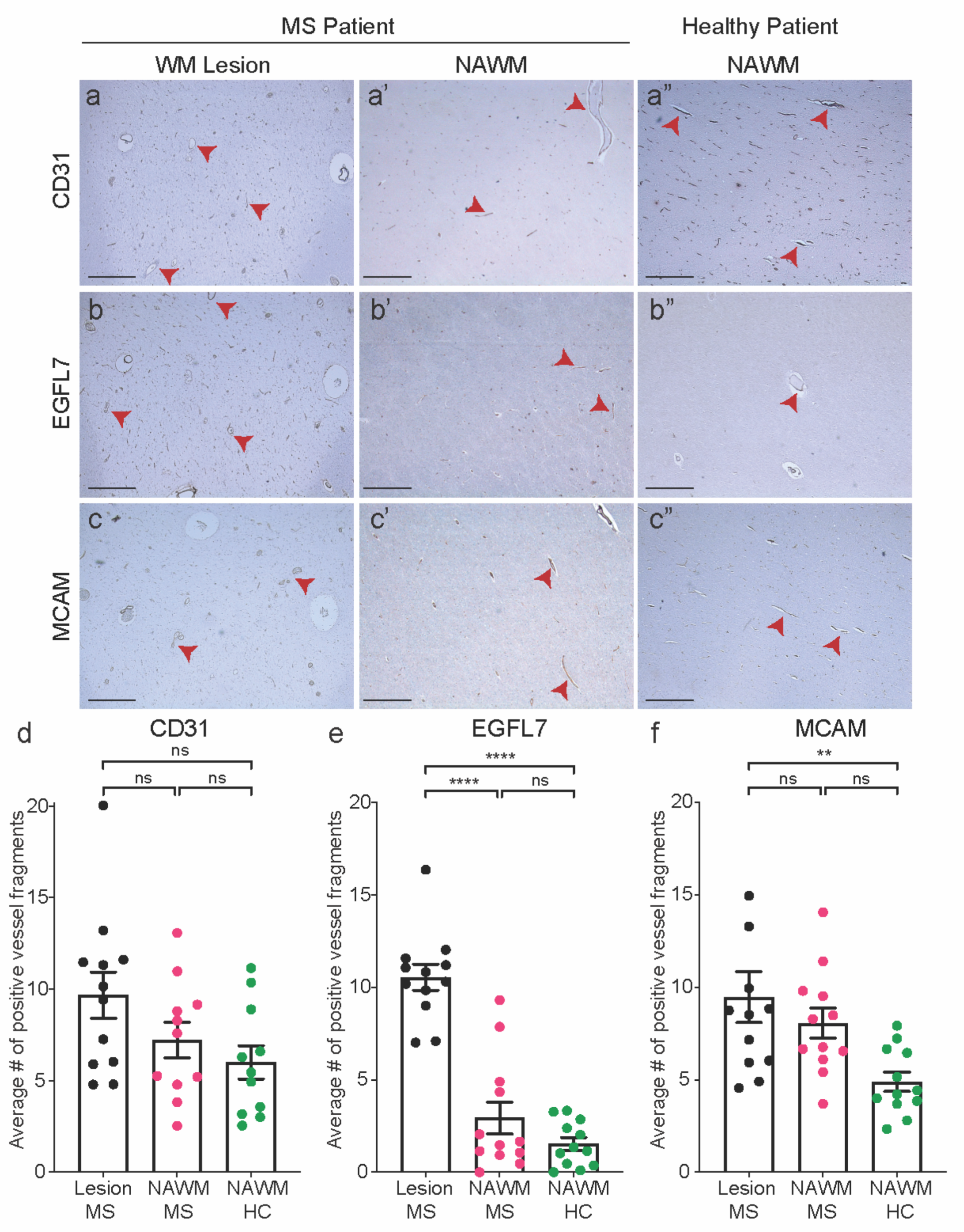
Increased expression of angiogenic markers in human multiple sclerosis lesions. **a, a’, a’’**) Immunohistochemistry (IHC) staining for CD31 (EC marker) in white matter (WM) lesions of MS patients (a), normal appearing white matter (NAWM) of an MS patient (a’) or a non-neurological control case (a’’). **b, b’, b’’**) IHC for the angiogenic marker, EGFL7, shows increased expression in WM lesions of MS patients (b) relative to NAWM of both the MS (b’) and a non-neurological control case (b’’). **c, c’, c’’**) IHC for the angiogenic marker, MCAM, shows increased expression in WM lesions of MS cases (c), relative to NAWM of non-neurological control case (c’’). **b-c’’**) Hematoxylin was used as a nuclear counterstain. **d, e, f**) Dotted bar graphs of the CD31^+^ average vessel density per given area (d), average EGFL7 (e), and MCAM (f) expression in blood vessels located in WM lesions and NAWM of MS patients (N = 11-12) and NAWM of non-neurological control patients (N = 11-12). Data represent mean ± SEM; n.s. = non-significant, *p < 0.05, **p < 0.01, ****p < 0.0001; two-tailed Student’s t test. Scale bars = 200 µm.

We asked whether transcriptomic features of pathological neo-angiogenesis are present in SPMS, characterized by progressive neurodegeneration and cumulative disability. We examined expression of neo-angiogenesis markers in SPMS cases using the spatial transcriptomics data from Kaufmann et al., 2022^86^. This study analyzed gene expression within an array of 100-μm diameter spots from MS lesions in SPMS and identified the presence of angiogenesis near MS lesions^86^. We binned all spots by their distance from MS lesion borders in each case, using bins of 0.5 mm width, and calculated the fraction of spots within each bin where neo-angiogenesis genes were detected. White matter (WM) spots near MS lesions showed higher expression for *SERPINE1 and MARKSL1* mRNAs, two regulators of angiogenesis^84,87^, compared to either gray matter (GM) area near the lesion, or a distant region (sulcus area; **Extended Data Figure 5e, g**). In contrast, *EGFL7* and *ECM1* mRNAs showed higher expression in the GM near the lesion (**Extended Data Figure 5d, f**). Thus, some molecular markers of pathological neo-angiogenesis are also present in SPMS.

### Treatment with the anti-VEGF-A monoclonal antibody r84 at the EAE onset downregulates transcriptomic features of pathological neo-angiogenesis

Upregulation of VEGF-A and TGF-β signaling in vECs from acute and chronic EAE together with features of neo-angiogenesis (**Figure 3a, c**) implicates their role as potential drivers of neo-angiogenesis in neuroinflammation. In addition to promoting angiogenesis, VEGF-A also induces vascular permeability by triggering the disassembly of adherens and tight junctions in endothelial cells through VEGFR2 phosphorylation and activation^88^. r84 is a fully human anti-VEGF-A monoclonal antibody that binds to both human and mouse VEGF-A and selectively blocks its interactions with VEGFR2 inhibiting tumor angiogenesis^43,44^. To test how effectively r84 inhibits VEGF-A-driven vascular permeability in mouse brain endothelial cells (mBEC), we measured trans-endothelial electrical resistance (TEER) over time (a readout of barrier permeability), in control mBEC and those treated with VEGF-A (100 ng/mL) and a 100-fold molar excess of either r84 or a human IgG isotype control. VEGF-A/IgG treatment induced a significant drop in TEER compared to control (no treatment); this effect was partially rescued by r84 addition (**Extended Data Figure 6a**). Bulk RNA sequencing of control, VEGF-A/IgG, and VEGF-A/r84-treated mBECs revealed significant changes in gene expression and signaling pathway activation as determined by the principal component analysis (PCA), differential expression analysis, and GSEA (**Extended Data Figure 6b; Extended Data Table 2**). VEGF-A increased expression of “pro-angiogenic” and “pro-inflammatory” gene signatures, but decreased expression of “oxidative phosphorylation” signatures; these effects were counteracted by r84 (**Extended Data Figure 6c**). However, r84 treatment did not reduce upregulation of “Wnt/β-catenin signaling” gene signatures driven by VEGF-A treatment in mBECs (**Extended Data Figure 6c**).

To test whether VEGF-A inhibition reduces transcriptomic features of neo-angiogenesis in EAE and affects clinical outcomes, we administered 5 mg/kg of either r84 or the human IgG1 isotype control to EAE mice every 3 days starting at 6 d.p.i. (before disease onset) until tissue collection at chronic stage (28 d.p.i) (**Figure 7a**). r84 treatment lowered the incidence of more severe EAE in mice relative to the IgG (**Figure 7b**), although there was no significant difference in clinical course of EAE, disease onset, peak clinical score and score on the day of collection between two treatments (**Figure 7c-e**). To determine whether r84 treatment suppresses the transcriptomic signatures of neo-angiogenesis in vECs during EAE, we isolated spinal cord ECs from r84- and IgG-treated chronic EAE mice (28 d.p.i.) mice for scRNA-seq analysis [n=3 r84- and 3 IgG-treated chronic EAE samples (6 and 8 mice, respectively)]. All samples were processed in the same batch (**Extended Data Table 1-1**). The clustering of spinal cord ECs isolated from both treatments and visualized with the t-SNE plots (**Figure 7f, g**) reflected the same AV zonation observed without treatment (**Figure 1c-e**) Venous EC (vECs and vnECs), but not aECs or cECs, from the r84-treated EAE mice formed separate clusters from IgG-treated ones (**Figure 7h**). GSEA of differentially expressed genes in r84-versus IgG-treated venous ECs from chronic EAE revealed suppression of gene signatures associated with “angiogenesis”, “tip cells”, “antigen presentation”, and “apoptosis”, while upregulation of those related to “BBB” and “canonical Wnt signaling” (**Figure 7i, Extended Data Table 1-10**). BBB genes related to “tight junctions”, “transporters”, “Wnt/β-catenin signaling”, and “transcytosis” were upregulated in vECs, whereas “angiogenic” signatures were downregulated (**Figure 7j, k**). Therefore, r84-treatment suppressed neo-angiogenic transcriptomic signatures in chronic EAE, similar to its effects *in vitro* (**Extended Data Figure 6b)**.

**Figure 7.**
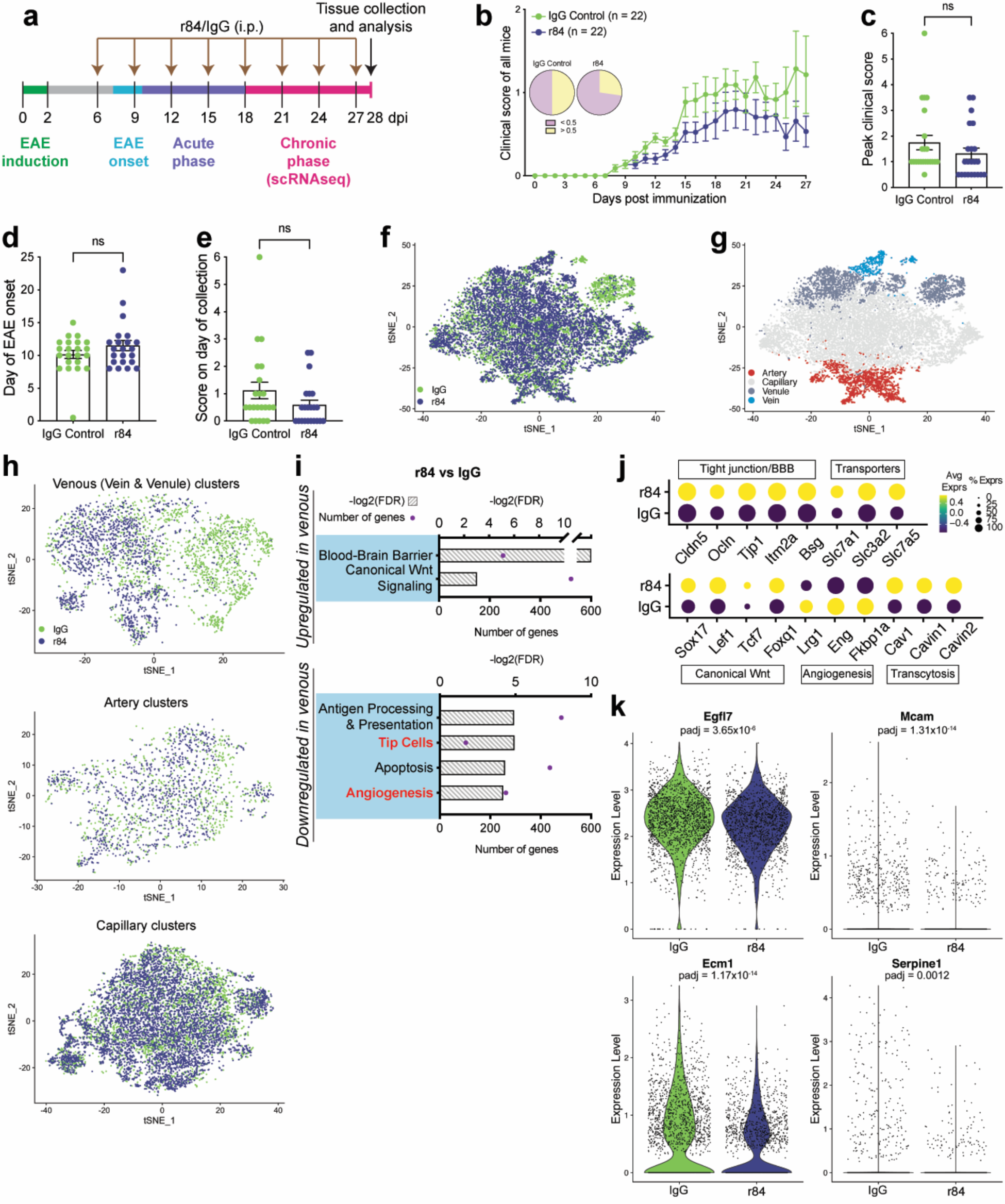
Gene set enrichment analysis (GSEA) reveals that transcriptome signatures of neo-angiogenesis are downregulated in spinal cord ECs of chronic EAE mice following treatment with r84. **a**) Diagram of the experimental procedure. Following EAE induction, mice were injected intraperitoneally (i. p.) every 3 days with either 5 mg/kg anti-VEGF monoclonal antibody r84, or an isotype control immunoglobulin G1 (IgG1), from 6 until 27 d.p.i. (chronic phase). Tissue was collected at 28 d.p.i. for analyses. **b**) Clinical EAE score curves for r84- and IgG1-treated mice. The inset pie charts depict the fraction of mice with clinical scores < 0.5 (pink) and > 0.5 (yellow) in the two treatment groups. The IgG1-treated EAE mice exhibit higher incidence of clinical scores > 0.5 compared to r84-treated EAE mice. **c-e**) Dotted bar graphs show no significant difference between the peak clinical score (c), day of disease onset (d), and score on the day of collection (e) between two groups. **f**) t-SNE visualization of ECs isolated from r84-treated (n = 3 samples; 6 mice) and IgG-treated (n = 3 samples; 8 mice) EAE spinal cords, color-coded for treatment. **g**) t-SNE plot, color-coded for EC subtype, shows that r84- and IgG treated ECs in this representation are visually organized along the arteriovenous zonation gradient. **h)** t-SNE plots, color coded for treatment, showing clustering of venous (vein & venule), arterial and capillary ECs in r84- and IgG-treated EAE mice. Only venous (vein & venule) ECs have non-overlapping clusters for r84 and IgG treatment groups. **i**) Pathways/processes significantly up- and downregulated in r84-treated relative to IgG-treated chronic (28 d.p.i) EAE venous (vein & venule) ECs. The gene ontology enrichment plots, derived using the GSEA algorithm, show that expression of neo-angiogenic gene sets and pathways (highlighted in red) are significantly downregulated in r84 EAE ECs. **j**) Dot plot shows significant upregulation of tight junction/BBB, transporter, canonical Wnt and transcytotic genes, and significant downregulation of angiogenic genes, in r84-treated venous (vein & venule) chronic (28 d.p.i) EAE ECs. **k**) Violin plots depicting reduction in *Egfl7*, *Mcam*, *Ecm1* and *Serpine1* expression levels in r84-relative to IgG-treated chronic (28 d.p.i) EAE venous (vein & venule) ECs. Data represent mean ± SEM; * p < 0.05, **p < 0.01, ****p < 0.0001; b) one-way ANOVA with repeated measures; c, d, e) two-tailed Student’s t-test.

To understand the effects of r84 treatment on the EAE vascular pathology, we examined whether neo-angiogenesis was reduced in r84-treated spinal cord in chronic EAE. We compared vEC proliferation and venous coverage in chronic EAE spinal cords (28 d.p.i) between r84- and IgG-treated groups. There was a significant reduction in vEC proliferation (Ki67^+^ Ecs), EphB4^+^ venous coverage and vein diameter in r84-compared to IgG-treated EAE spinal cords (**Figure 8a-c, g**). Similarly, the vascular coverage of Endomucin^+^ vECs was significantly reduced in EAE mice after r84 treatment **(Figure 8d, e)**. Consistent with scRNA-seq results (**Figure 7**), these data demonstrate that neo-angiogenesis (vEC proliferation, increased venous diameter and coverage) observed in EAE was reduced by inhibiting VEGF-A signaling with r84. In contrast, the area of serum IgG and fibrinogen leakage across the BBB in chronic EAE spinal cords were not significantly different between the two groups **(Figure 8d, f; Extended Data Figure 7b, h)**. Thus, although r84 administration reduced neo-angiogenesis in chronic EAE, it did not rescue increased BBB permeability, a critical pathological feature of the vasculature in EAE/MS.

**Figure 8.**
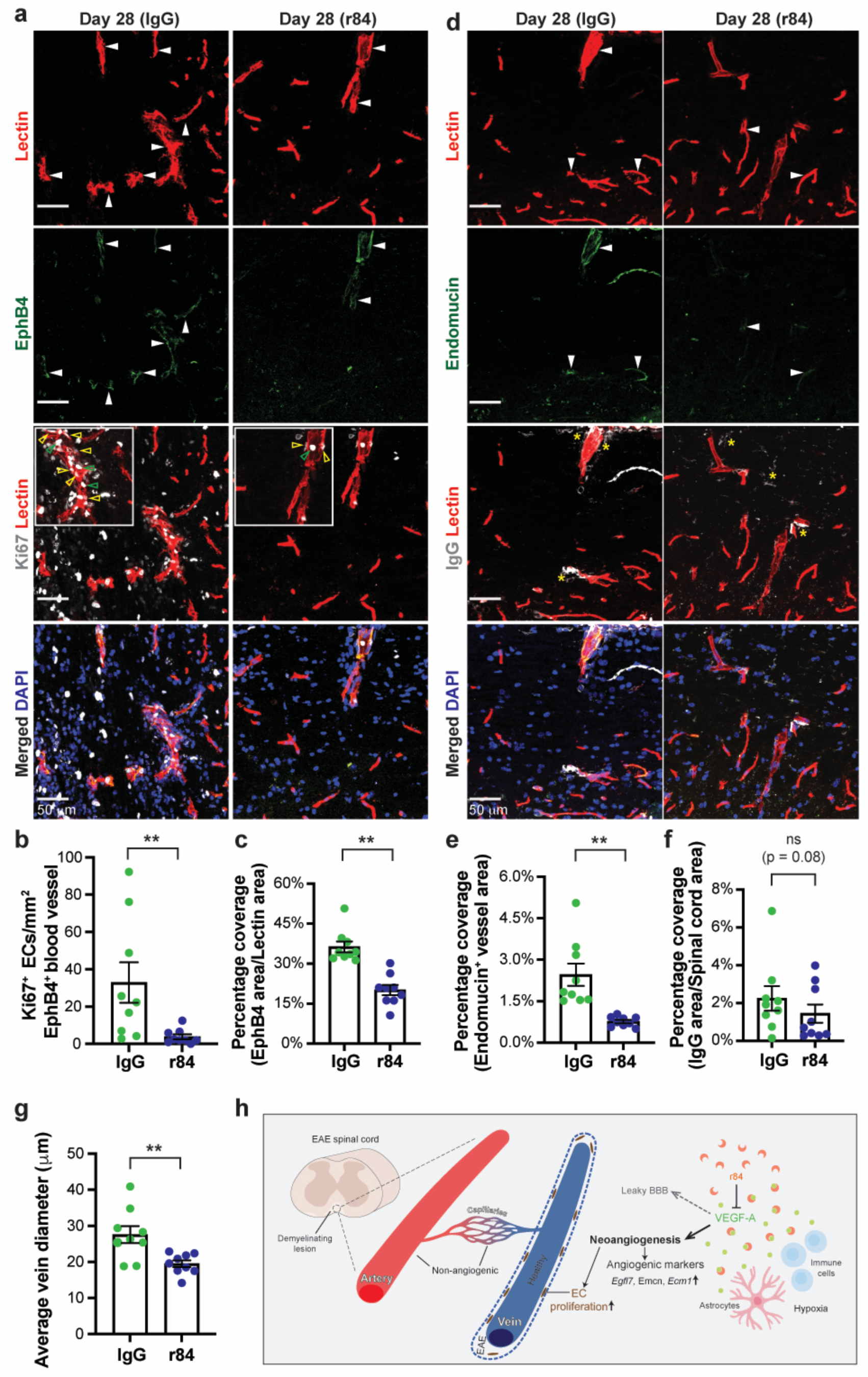
r84 treatment reduces pathological neo-angiogenesis in vein ECs in EAE, but does not rescue BBB leakage. **a)** Sagittal sections of IgG- and r84-treated 28 d.p.i. EAE spinal cords were immunostained with antibodies against EphB4, Ki67 as well as DAPI following cardiac perfusion with Lectin (marks perfused ECs). White arrowheads point at EphB4^+^/Lectin^+^ (double positive) vessels. Empty yellow arrowheads point at Ki67^+^ ECs. Empty green arrowheads point at Ki67^+^ immune cells. **b, c)** Dotted bar graphs of the number of Ki67^+^ ECs in EphB4+ vessels (b) and EphB4^+^ vascular area (c) show lower EC proliferation and venous coverage in r84-treated spinal cords compared to the control (n = 9 mice /group). **d)** Sagittal sections of IgG- and r84-treated 28 d.p.i. EAE spinal cords were immunostained with antibodies against Endomucin (marks veins), endogenous serum IgG and DAPI after cardiac perfusion with Lectin. White arrowheads point at Endomucin^+^/Lectin^+^ (double positive) vessels. Yellow asterisks point at serum IgG leakage around blood vessels. **e-f**) Dotted bar graphs show quantification of Endomucin^+^ vascular area (e) and IgG leakage area (f). There was significantly lower venous coverage but no significant difference in IgG leakage area in r84-treated spinal cords compared to the control (n = 9 mice / group). **g)** Dotted bar graph showing quantification venous diameter reveals significant reduction in r84-treated spinal cords compared to the control (n = 9 mice / group). **h)** Model illustrating the putative mechanism of pathological neo-angiogenesis in EAE. Neuroinflammation induces enlargement of veins and venules (via vEC proliferation), with leaky BBB, via neo-angiogenesis in the white matter lesions of the spinal cord, where there is a significant presence of infiltrating immune cells. Hypoxic conditions induce secretion of VEGF-A from either infiltrating immune cells (e.g. macrophages), or CNS resident cells (e.g. astrocytes), resulting in activation of VEGF-A signaling pathway in vECs. Blockade of VEGF-A signaling through administration of r84 reduces neo-angiogenesis. But r84 treatment does not reduce disease severity, immune cell infiltration and BBB permeability. Our study identifies VEGF-A signaling as a driver of neo-angiogenesis in neuroinflammation.

To understand why r84 treatment did not affect EAE clinical outcome, we examined the number of lesions, extent of demyelination and immune infiltration in r84- and IgG-treated EAE spinal cords. While EC proliferation and venous coverage were downregulated significantly in r84-treated EAE mice (**Figure 8a-c**, **Extended Data Figure 7c**), the number of lesions, the percentage of demyelinated WM area (measured by the absence in fluoromyelin staining), and the number of infiltrating CD4^+^ T helper cells and the CD68^+^ infiltrating macrophages, were not significantly different between the two treatment groups (**Extended Data Figure 7a-b, d-g**). Overall, our findings indicate that VEGF-A signaling drives pathological neo-angiogenesis in EAE from vECs. Although its blockade reduces gene signatures and histopathological features of neo-angiogenesis, it is not sufficient to rescue BBB leakage, and consequently immune cell infiltration into the CNS, demyelination and EAE clinical outcomes.

## DISCUSSION

The presence of neo-angiogenesis and high levels of VEGF-A has been documented in both MS and EAE for several decades^20–23,27,32^. Previous functional studies using monoclonal antibodies (Bevacizumab, B20.4-1-1) to block VEGF-A signaling have concluded that VEGF-A-driven neo-angiogenesis worsens EAE clinical outcomes^26,28^. However, the timing and relation of neo-angiogenesis to EAE/MS progression, its molecular characterization, and the vascular origin and signaling pathways driving neo-angiogenesis in neuroinflammation are poorly understood. Here, we employed scRNA-seq and GSEA, together with fluorescent *in situ* hybridization and immunofluorescence validation to demonstrate that venous ECs, primarily in acute EAE, express transcriptomic signatures of neo-angiogenesis together with activation of pro-angiogenic VEGF-A and TGF-β signaling (**Figure 3**). Thus, venous ECs are the primary EC subtype responsible for pathological angiogenesis in neuroinflammation (**Figure 8h**). Several neo-angiogenesis markers, identified through scRNA-seq in EAE, are also upregulated in human MS lesions from both RRMS and SPMS cases. This upregulation is accompanied by a moderate, albeit not significant, increase in vascular density (**Figure 6; Extended Data Figure 5**). Through functional validation studies, we show that treatment with the VEGF-A blocking antibody r84^43,44^ at the disease onset reduces neo-angiogenic transcriptomic signatures and EC proliferation *in vivo*, but does not restore BBB function, or significantly ameliorate EAE pathology. Below, we discuss a putative model for pathological angiogenesis in neuroinflammation and therapeutic implications for MS.

Prior studies employing bulk or scRNA-seq of the endothelium in EAE have shown that ECs acquire transcriptomic features of inflammation and lose expression of those associated with the BBB^50,51^. Through scRNA-seq and *in vivo* validation, we demonstrate for the first time that transcriptomic signatures and features of angiogenesis arise from vein/venule ECs in neuroinflammation. This was achieved through isolation of sufficient ECs from each vessel subtype from healthy, CFA controls and disease states to identify gene signatures and pathways that were up- and downregulated in neuroinflammation. Since the number of angiogenic ECs arising from vEC/vnECs is small, they were likely missed in previous analyses^50,51^. Nevertheless, the identification of vECs as the source of neuroinflammation-related angiogenesis is somewhat surprising. The classical model suggests that angiogenesis occurs from pre-existing capillaries, whereby cECs respond to a hypoxia-induced VEGF-A gradient and differentiate into migratory tip and proliferating stalk cells that expand the vascular network (reviewed in ^89,90^). Recent studies have challenged this dogma. Coelho-Santos *et al*. have shown that CNS angiogenesis in the neonatal mouse brain is initiated from cortical ascending venules^72^. Lee *et al*. have utilized EC vascular subtype-specific inducible reporter mice to fate map ECs in the developing postnatal retinal vasculature, and discovered that ∼80% of ECs are of venous origin at both P7 and P25^74^. Moreover, neo-angiogenic sprouts arise exclusively from veins in the oxygen-induced retinopathy (OIR) model^74^. Rohlenova *et al.* have employed scRNA-seq and trajectory inference to predict that vECs differentiate into neo-angiogenic tufts during choroidal neovascularization (CNV), a model for wet age-related macular degeneration (AMD)^91^. Their data also suggests that disease-specific EC population, which contains clusters of proliferating and tip ECs, exhibit vein marker expression^91^. Lastly, a recent scRNA-seq study of the lung tumor vasculature has also found that tumors have a higher proportion of vECs, compared to the normal vasculature^63^.

Why are vein/venule, but not arterial/capillary ECs angiogenic in neuroinflammation? Postcapillary venules and veins are the primary sites of immune cell infiltration and BBB damage in MS/EAE^1,2^. Activated T cells that infiltrate into the hypoxic CNS parenchyma secrete VEGF-A^92,93^, which can promote angiogenesis locally (**Figure 8h**). Macrophages, which make up a large proportion of cellular infiltrates in MS/EAE also secrete numerous pro-angiogenic factors, including VEGF-A, FGF, PDGF and Angiopoetin-1 and -2^94^ (**Figure 8h**). They also produce the matrix metalloproteinases MMP-7 and -9^95^, which can digest the ECM surrounding the veins/venules to free ECM-bound VEGF-A, while simultaneously creating a path through the vascular basement membrane for EC migration. Another explanation for the venous origin of neo-angiogenesis in neuroinflammation could be related to differences in shear stress levels present in arteries, capillaries and veins which regulates phosphorylation of VEGFR2, VE-Cadherin, CD31 and Erk1/2 (reviewed in^96^). High shear stress in arteries may obstruct EC proliferation and migration, whereas low shear stress in veins may be conducive to neo-angiogenesis.

Our scRNA-seq analyses of spinal cords ECs from control and EAE mice revealed that gene sets associated with VEGF-A, but not Wnt/β-catenin, signaling and neo-angiogenesis are specifically enriched in vECs in neuroinflammation, implicating VEGF-A as a possible driver of neo-angiogenesis in EAE (**Figure 8h**). Previous studies have shown that administration of VEGF-A inhibitors such as bevacizumab ^26^, B20-4.1.1^28^ or SU5416^24^ at the EAE induction improve clinical outcomes in EAE mice. It has been unclear, however, if these VEGF-A inhibitors exert their therapeutic effects directly on CNS angiogenesis in neuroinflammation, or indirectly by dampening the peripheral immune response resulting in less CNS immune infiltration. Treatment with SU5416 (Semaxinib), a potent and selective inhibitor of VEGFR2, reduces immune filtration in the CNS^24^. Similarly, Macmillan *et al.*, have shown that prophylactic administration of bevacizumab (humanized monoclonal antibody) and B20-4.1.1 (monoclonal antibody that binds both human and mouse VEGF-A) prior and throughout EAE onset in mice reduces peripheral T cell responses, decreases CD4^+^ T cell recruitment into the spinal cord and IL-17A and IFNγ secretion in response to *in vitro* stimulation with MOG_35-55_ ^26,28^. However, bevacizumab does not bind murine VEGF-A, and therefore it is puzzling how it improves EAE clinical outcomes^26^. In our study, we used r84, a humanized anti-VEGF-A monoclonal antibody that binds both human and mouse VEGF-A and selectively blocks its interactions with VEGFR2 receptor ^43,44^ to test the contribution of VEGF-A-mediated neo-angiogenesis in EAE pathology. Our data demonstrate that r84 administration prior to EAE onset reduces transcriptomic signatures and pathological features of neo-angiogenesis including vEC proliferation, increased venous diameter and coverage observed in EAE (**Figure 7**). Although r84 can inhibit acute VEGF-A driven endothelial barrier breakdown *in vitro* **(Extended Data Figure 6a)**, BBB leakage in chronic EAE was not improved by r84 administration **(Figure 8d, f; Extended Data Figure 7b, h)**. This could be due to difference between *in vitro* and *in vivo* inflammatory environments or timing (acute vs chronic). Since increased BBB permeability is a critical pathological feature of EAE that permits immune cell infiltration into the CNS and pathology, there was no significant reduction in T cell or macrophage infiltration, demyelination and EAE clinical outcomes upon r84 administration. Our findings contrast with previously published studies showing that VEGF-A inhibitors such as bevacizumab ^26^, B20-4.1.1^28^ or SU5416^24^ can improve clinical outcomes in EAE mice. The B20-4.1.1 treatment regimen in previous studies may have inhibited VEGF-A binding to VEGFR1 and/or soluble VEGFR1 (Flt1/sFlt1), which could affect the amount of VEGF-A available to bind to VEGFR2. This may explain, in part, the difference in clinical outcomes between previous B20-4.1.1 and our r84 treament regimes in EAE. Further in-depth analyses are warranted to identify the key mechanistic differences between these two treatments. Although it is likely that the beneficial effects of bevacizumab in EAE are off-target since it does not bind murine VEGF-A, other VEGF-A inhibitors, such as B20-4.1.1, likely dampen the initial priming of T-cell mediated immunity critical for EAE initiation, since they were administered at the time of EAE induction. Since we have administered r84 at the time of disease onset (day 6), bypassing the initial peripheral immune activation, it is possible that anti-angiogenic therapies in combination with immunomodulatory therapies may benefit MS progression. Further studies are warranted to combine r84 treatment with immunosppressive therapies (that do not affect neo-angiogenesis) to ascertain the effect of this combination therapy in EAE clinical outcome.

In addition to its role in angiogenesis, VEGF-A is a key regulator of vascular permeability through multiple mechanisms (reviewed in^88^). Activation of VEGF signaling downregulates expression of Claudin-5 and Occludin^97^, induces phosphorylation of Occludin and ZO-1^98,99^ and disrupts ZO-1 and Occludin organization^100^, resulting in TJ disassembly in brain microvascular endothelial cells (BMECs). VEGF-A also promotes VE-cadherin endocytosis and barrier dysfunction^101^. DE analysis revealed that expression of the TJ proteins, Claudin-5, Occludin and ZO-1 (*Tjp1*), were significantly upregulated in vein Ecs after r84 treatment, likely due to inhibition of VEGF-A signaling^102^. Therefore, r84-mediated VEGF-A inhibition in EAE may reduce both vEC neo-angiogenesis and promote restoration of several BBB properties (tight junctions, Wnt/β-catenin signaling, transporters) at the transcriptome level (**Extended Data Fig. 5**). However, we show this is not sufficient to restore BBB function, likely due to contribution of other factors (e.g. inflammatory cytokines) in chronic EAE^103^.

In conclusion, our study suggests a model in which VEGF-A derived from either reactive astrocytes^104^ or infiltrating immune cells induces neo-angiogenesis from vein ECs in EAE/MS through upregulation of several angiogenic markers such as Egfl7, Emcn, Ecm1, Serpine1 and Mcam which promotes venous EC proliferation, vein diameter expansion and increased vein coverage (**Figure 8h**). Inhibition of VEGF-A signaling with r84 reduces transcriptomic features of neo-angiogenesis (including downregulation of Egfl7, Mcam, Ecm1, Serpine1 and Emcn), without affecting BBB leakage, immune cell activation or their CNS infiltration and EAE clinical outcomes (**Figure 8h**). Several markers of pathogenic angiogenesis identified through our scRNA-seq in EAE are also upregulated in either acute or chronic demyelinating MS plaques in RRMS or in SPMS cases, through *in situ* hybridization analysis and post-hoc spatial transcriptomic analysis of published data^86^, respectively. SPMS has no current treatment. Our results advocate for VEGF-A inhibition as a possible therapeutic intervention in the early phases of RRMS to reduce neo-angiogenesis, which may improve long-term neurological deficits in patients as they transition to SPMS.

## Online Methods

### Mice

All experiments were approved by the Institutional Animal Care and Use Committees (IACUC) at Columbia University Irving Medical Center. Animals used for breeding and EAE induction were housed in sex-matched cages of up to five mice in the vivarium with access to food and water *ad libitum*. Wild-type C57BL/6J, obtained from the Jackson Laboratory (Maine), were used for EAE experiments to collect spinal cords from CFA control, acute and chronic EAE for analysis of neo-angiogenesis via immunofluorescence and RNA fluorescent in situ hybridization (FISH). TCF/LEF::H2B::eGFP (Wnt reporter) mice^9,42^, purchased from Jackson Laboratory, were used for single-cell RNA sequencing (scRNA-seq) of control and EAE spinal cord endothelial cells (ECs), to track Wnt/β-catenin pathway activation in ECs *post hoc* within the RNA-seq data. The mice were genotyped for GFP with primers [CCCTGAAGTTCATCTGCACCAC (forward); TTCTCGTTGGGGTCTTTGCTC (reverse)] as described^42^.

### Induction of MOG_35-55_ EAE

The induction of MOG_35-55_ EAE in mice has been described^9,42^. Briefly, 10–12-week-old female mice were immunized by subcutaneous injection with 100 µg of myelin oligodendrocyte glycoprotein 35-55 (MEVGWYRSPFSRVVHLYRNGK-COOH) in complete Freund’s adjuvant (CFA) containing 200 µg heat killed mycobacterium tuberculosis H37Ra (Difco). 400 ng of *Bordetella pertussis* toxin was administered intravenously at day 0 and day 2. Mice were examined for clinical EAE signs starting at 5 days post immunization (d.p.i) using the scale (0.5-point gradations for intermediate presentation): 0, no signs; 1, flaccid tail; 2, hindlimb paresis; 3, hindlimb paralysis; 4, hindlimb and forelimb paralysis; 5, moribund; 6, death. Control animals immunized with CFA emulsion and pertussis toxin did not exhibit any clinical signs. Spinal cord tissue was collected at the peak acute (16 d.p.i), the beginning of chronic (28 d.p.i), or chronic (40 d.p.i) EAE, as well as from healthy and CFA controls, for scRNAseq, immunofluorescence and RNA *in situ* hybridization analyses of angiogenic marker expression.

### Injection of VEGF-A monoclonal antibodies in EAE

Mice induced with MOG_35-55_ EAE were placed into two groups at 6 d.p.i.: Group 1 (r84-treated) was injected intraperitoneally with 5 mg/kg of the anti-VEGF-A monoclonal antibody, r84^43^ every 3 days until tissue collection at 28 d.p.i. Group 2 (IgG-treated) was injected with 5 mg/kg of the immunoglobulin G isotype control ([IgG] Biolegend, Cat. # 403502) every 3 days until spinal cord tissue collection at 28 d.p.i. (**Figure 7a**), for analysis of angiogenesis, immune infiltration and leakage, demyelination and scRNA-seq.

### Isolation of endothelial cells from control and EAE spinal cords

Mice were anesthetized with isoflurane and perfused for 4 minutes with ice-cold sterile PBS. The spinal cord was dissected into Earl’s balanced salt solution (EBSS) and dissociated into a single cell suspension using an established protocol^105^. The single cell suspension was resuspended in 200 µl of CD16/CD32 Fc block (BD Biosciences, Cat. # 553141; 1/200 in FACS buffer) and incubated at room temperature for 15 minutes. Samples were washed with 2 ml of FACS buffer, and 100 µl were kept aside for unstained and fluorescence minus one (FMO) controls. The cells were resuspended in 200 µl of anti-CD31-APC (Biolegend, Cat. # 102410; 1/200 in FACS buffer) antibody to label endothelial cells (ECs) and incubated on ice in the dark for 30-60 mins. Following antibody labeling, the samples were washed twice with 2 ml of FACS buffer and resuspended in 400 µl of propidium iodide (Thermofisher, Cat. # P1304MP; 1/10,000 in FACS buffer). Gates for the surface stain were set using the unstained and FMO controls. The CD31-APC positive ECs were sorted out using the BD FACS Aria Flow Cytometer (Columbia Stem Cell Initiative Flow Cytometry Core) and processed for scRNA-seq.

### mBEC culture and TEER measurements

Primary mouse brain endothelial cells (mBECs; Cell Biologics C57-6023) were grown to confluency on Poly-D-Lysine (10ug/mL) and Type IV collagen (20ug/mL) coated 96W20idf PET plates (Applied Biophysics), for TEER experiments and 10 cm plates for bulk RNA-seq with complete mouse endothelial cell medium kit (Cell Biologics M1168) and 5% FBS. TEER was measured in real time using an electric cell-substrate impedance sensing (ECIS) instrument (Applied BioPhysics). Once confluency was reached, cells were switched to 1% FBS medium without growth factor supplements. After 24 hours in low serum, cells were treated with control medium, VEGF-A (100ng/mL, Cat #493-MV, Lot #RQ0321011) and Human IgG1 (Bioxcell, Cat #BE0297, Lot #786222A2), or VEGF-A (100ng/mL) and r84 (generated by R. A. Brekken). IgG and r84 were used in 100-fold molar excess as reported^43^.

TEER was measured for 48 hours after treatment (96 hours total). The change in TEER was determined by calculating the average resistance values during a 1-hour period following treatment (acute) and the last 1-hour period of recording (chronic), and then subtracting this value from the average resistance of the well during the 1-hour period prior to treatment. Values were averaged across 8 wells for each condition to generate time course and conditions were compared by unpaired two-tailed Student’s t-test. Cells were also collected at 48 hours after treatment and RNA was extracted using Rneasy Mini Kit (QIAGEN) for bulk RNA sequencing.

### Single-cell RNA sequencing analysis

**Sample Size:** For scRNA-seq, we used two mice per sample to collect sufficient ECs from spinal cords. The sample size was: a) healthy mice: n = 3 samples (n = 6 mice) from which we isolated 4251, 2907, 4531 ECs from each sample; b) CFA disease control: n = 3 samples (n = 6 mice) from which we isolated 3377, 4360, 3595 ECs per sample; c) acute EAE: n = 4 samples (n = 8 mice) from which we isolated 2176, 1861, 2002, 1929 ECs per sample; and d) chronic EAE: n = 4 samples (n=8 mice) from which we isolated 3110, 2381, 4603, 4226 ECs per sample. For scRNA-seq of r84- and IgG-treated EAE mice we used 2-4 mice per sample to collect sufficient ECs from spinal cord (IgG: n = 3 samples (8 mice) from which we isolated 2381, 1429 and 2374 ECs per sample; r84: n = 3 samples (6 mice) from which we isolated 921, 3456 and 3987 ECs per sample) (**Extended Data Table 1-1**).

The 10x Genomics Chromium^106,107^ platform was used to profile cells (at a median of ∼2000 genes/cell), and sequencing data was processed using the Cellranger v3.0 analysis pipeline to align reads and generate feature-barcode matrices. The Seurat R package^45^ was used to read the output of the Cell Ranger pipeline and merge data from all the samples (controls and EAE) into a single R object. The standard pre-processing workflow for scRNAseq data was carried out in Seurat, which consisted of the selection and filtration of cells based on quality control metrics (number of unique genes detected in each cell > 200, total number of molecules detected within a cell > 1,000 and < 50,000, percentage of reads within a cell that map to the mitochondrial genome < 20), data normalization and scaling, and the detection of highly variable features. Next, linear dimensionality reduction (PCA) was performed on the scaled data using the previously determined highly variable features (2,000 genes), an alternative heuristic method (’Elbow plot’) was implemented to determine the ’dimensionality’ of the dataset (resulting in 30 PCs being retained), and cells were clustered by applying the weighted shared nearest-neighbor graph-based clustering method^108^ using the first 30 PCs and resolution value 1.5. Lastly, the 30 PCA dimensions were further reduced into two-dimensional space using the non-linear dimensional reduction technique, t-distributed stochastic neighbor embedding (t-SNE), to visualize and explore the scRNA-seq dataset and each of its unique EC clusters. This analysis workflow produced 23 EC clusters. These clusters are not necessarily biologically distinct, as the main purpose of the ’over-clustering’ step was to establish clear boundaries between cells belonging to each of the major vessel subtypes (arteries, capillaries, venules, veins) using EC subtype-specific marker gene expression. Cluster-specific expression of these EC subtype-specific genes were then used to assign each of the 23 clusters to a major vessel subtype. This approach based on over-clustering and then re-merging based on computational or biological meaning avoids pitfalls arising from incorrect clustering boundaries with lower resolution^47^.

### Differential Gene Expression Analysis

The Seurat “FindMarkers” function with the default Wilcoxon Rank Sum test was implemented to compile the lists of differentially expressed genes, by comparing expression profiles of disease enriched (acute and chronic) EC clusters to control EC clusters, and r84-treated vEC clusters to IgG-treated vEC clusters from chronic EAE.

### Gene Set Enrichment Analysis (GSEA)

Following DE analysis, a score was calculated for each gene using the formula: Gene score = -log_10_(pval) x sign (log_2_fc). Up- and downregulated genes had a positive and negative score, respectively. The numeric value of the gene score was inversely correlated with the degree of significance, that is, genes that had the highest statistically significant difference in expression and the lowest p-values had the highest score, while non-significant genes with high p-values had a correspondingly low gene score. This score was used to rank each gene in DE gene list (**Extended Data Table 1-3 to 1-10**). New gene sets relevant to pathways, processes and cell types of interest (VEGF-A signaling, canonical Wnt/β-catenin signaling, non-canonical Wnt signaling, chemokine/cytokine signaling, TGF-β signaling, extracellular matrix, cell-cell adhesion, transporters, Notch signaling, angiogenesis, endothelial cell proliferation, endothelial cell migration, apoptosis, antigen processing and presentation, blood-brain barrier, inflammation, endothelial-to-mesenchymal transition, tip cells) were compiled as part of the overall gene set enrichment analysis (**Extended Data Table 1-2)**, using the GO, KEGG, DAVID and GSEA databases as published before^109–111^. The ranked list of DE genes, along with the compiled gene sets, were loaded onto GSEA 4.3.2 software, and run through ‘GseaPreranked’ with the following settings: Number of permutations = 1,000; Collapse/Remap to gene symbols = No_collapse; Max size: exclude larger sets = 1,500; all other settings were left at ‘default’. Gene sets with FDR q-value ≤ 0.10 were deemed significantly enriched (na_pos) or depleted (na_neg) in the cluster of interest (e.g., acute, or chronic EAE EC enriched cluster) relative to its counterpart (e.g., control EC enriched cluster) in the DE analysis, and plotted in the GSEA graphs. Although less rigorous than the standard p < 0.05 threshold, the 0.1 threshold is nonetheless the default in GSEA for standard enrichment analyses, so we maintained it here^52^.

### Bulk RNA sequencing

Bulk RNA-seq was performed by JP Sulzberger Columbia Genome Center. mRNA was enriched from total RNA samples using poly-A pull down and cDNA libraries were prepared using Illumina Truseq chemistry, followed by sequencing using Illumina NovaSeq 6000. Pseudoalignment of RNAseq reads was performed to a kallisto index created from transcriptome Ensembl v96, Mouse:GRCm38.p6 using kallisto (0.44.0). Differential gene expression analysis between control, VEGF-A/IgG, and VEGF-A/r84-treated samples was performed using DESeq2. Gene set enrichment analysis was performed using endothelial cell curated and hallmark gene sets. For heat map visualization, size factors were calculated using a median ratio method to normalize raw gene counts across samples (to adjust for differences in sequencing depth and library size) and z-scores were calculated for each gene’s normalized counts. To further assess overall similarity between samples, regularized logarithm transformation was performed on differentially expressed genes and principal component analysis was performed on transformed expression data.

### Immunofluorescence analysis

Mice used for immunofluorescence staining were anesthetized with isoflurane and perfused with PBS for 4 minutes followed by 4% PFA for 6 minutes. Spinal cords were extracted and post-fixed for 2 hours, washed three times in PBS for 30 minutes, then cryoprotected in 30% sucrose overnight and embedded in Tissue Plus O.C.T. Compound (Fisher Scientific, Cat # 4585). The tissue blocks were stored at -80 °C until use. Tissues were then sectioned at 12 µm on a Leica cryostat and stored at -80 °C until use. For staining, slides were air-dried for 20 minutes then placed horizontally in a humidified chamber. Slides were washed once in PBS for 10 minutes to remove excess OCT, then blocked for 1 hour in blocking buffer (10% BSA, 0.1% Triton-X-100 in 1x PBS). Primary antibodies (**Extended Data Table 3**) were diluted in 0.1% Triton-X-100 in PBS with 1% BSA. Primary antibody solutions were incubated overnight at 4 °C. Slides were washed three times for ten minutes with PBST (0.1% Triton-X-100 in PBS), then primary antibodies were targeted using associated fluorescently tagged secondary antibodies (1:1000 for Alexa488 and Alexa594, 1:500 for Alexa647). Secondary antibodies were diluted in PBST and incubated for two hours at room temperature. Nuclei were then labeled with DAPI (4′,6-diamidino-2-phenylindole, 1:20,000 in PBST) for 10 minutes, then slides were washed twice with PBST and twice with PBS. Slides were mounted with Vectashield (Vector Labs, Cat # H-1000), coverslipped and sealed with clear nail polish, and stored at -20 °C. Imaging of brain sections were performed with either a Zeiss LSM700 confocal microscope or a Zeiss AxioImager.

### Chromogenic immunohistochemistry of human tissue

Formalin-fixed human brain samples were obtained from the NIH Brain & Tissue Repository-California, Human Brain & Spinal Fluid Resource Center at the Veterans Affairs Hospital West Los Angeles Healthcare Center, Los Angeles, California, which is supported in part by National Institutes of Health and the US Department of Veterans Affairs under the CUIMC IRB protocol # AAAQ7343. Brain tissues from 16 MS and 16 non-neurological control patients were used in this study (**Extended Data Table 4**). Upon receipt, the fixed samples were washed with cold PBS then dehydrated in 30% sucrose before embedding in Tissue-Tek O.C.T. Compound (Sakura; Torrance, CA) and cryosectioned 12 µm thick. Single target immune-peroxidase staining was performed as described^42^. Hematoxylin counterstain was used for all stains except for CD31. For 2-target immune-peroxidase staining, targeting both CD31 and Ki67, the ImmPRESS Duet Double Staining Polymer kit (Vector Labs, Cat. # MP-7724, Burlingame, CA) was used, according to manufacturer’s recommendations. No nuclear counterstain was used in these experiments. All primary antibodies were diluted in 1% bovine serum albumin, 0.1% Triton-X, in PBS. Images were acquired with a Zeiss Axioimager using a color camera and 10X objective.

### *In situ* hybridization

Digoxigenin-labeled probes were synthesized from plasmids obtained from TransOMIC using the Roche Applied Science *in vitro* transcription kit (Sigma, 11175025910). The probes used for these experiments are listed in **Extended Data Table 5**. Mice used for *in situ* hybridization experiments were perfused with PBS only, brains and spinal cords were dissected out, embedded in OCT and frozen on dry ice, then stored at -80 °C. Sections were cut onto glass slides using a cryostat. Alkaline Phosphatase *in situ* hybridizations and fluorescent *in situ* hybridization (FISH) combined with immunofluorescence for vascular markers were performed as previously described^42,112^. Slides were imaged either on a Zeiss AxioImager, or an Zeiss LSM700 confocal microscope.

### Image analysis

Images were prepared and analyzed using ImageJ/FIJI software. For fluorescence measurements, at least three sections were imaged per sample, and in each case, multiple (8-12) representative regions of interest (ROIs) were acquired using the confocal microscope (20x objective). For each ROI, multiple Z-stacks were acquired, and maximal projection was performed in ImageJ to properly capture the full depth of the CNS blood vessels. Background correction was performed in ImageJ when required. For statistical analysis, the mean value for each animal was calculated as the mean signal from all acquired ROIs.

#### Analysis of vein EC proliferation and vein density

Spinal cord tissue sections were stained with the Ki67 (proliferation marker), EphB4 (vein and venule marker), along with a vascular marker (Caveolin-1/Lectin) and DAPI (**Extended Data Table 4**). Ki67^+^ venous nuclei were counted and normalized to the total venous area to get a readout of EC proliferation rate. The percentage area coverage (readout of vein density) by EphB4+ or Endomucin+ veins and venules over a vascular marker (Caveolin-1/Lectin) was quantified using ImageJ/FIJI.

#### Analysis of angiogenic marker expression

Spinal cord tissue sections were analyzed for the expression of angiogenic markers (Egfl7, Mcam, Ecm1 and Serpine1) and vessel/vein marker (Caveolin-1/Vcam1) with either FISH combined with immunofluorescence or immunofluorescence staining. The vessel area was outlined by signal thresholding using the ImageJ/FIJI software, and the mean fluorescence intensity (M.F.I.) of angiogenic marker expression within the vasculature was measured using the histogram analysis feature.

### Statistical analyses

Statistical analyses of EAE disease course and parameters (day of disease onset, peak disease score and score on the day of collection), vascular subtype cell proportions within the scRNAseq data, vein EC proliferation and vein density, angiogenic marker expression and flow cytometry data were performed using GraphPad Prism. Gene set enrichment analysis was performed using the GSEA v4.3.2 Mac App. Data were analyzed by Student’s t-test, one-way, two-way, or repeated measures (RM) ANOVA followed by Sidak’s, Tukey’s, or Bonferroni’s post-hoc corrections. Significance (α) was set at 0.05. Data in the dotted bar graphs are represented as mean ± SEM, where each dot is the mean value from each animal.

## Data availability

The raw and analyzed mouse endothelial scRNA-seq and bulk RNA-seq datasets supporting this manuscript are archived at the NIH GEO repository under accession number GSE210776.

## Acknowledgements

We thank Jeroen Bastiaans and Heidi Stuhlmann at Weill Cornell Medical College, New York for providing the mouse Egfl7 mRNA probe, and help optimizing the staining of the human tissue with the EGFL7 protein, Tyler Cutforth for advice with the preparation of mRNA probes and FISH experiments, Michael Kissner at the Columbia Stem Cell Initiative Flow Cytometry Core and Erin Bush at the Columbia Single-Cell RNA-sequencing facility for their advice and help with FACS isolation and single-cell RNA-sequencing experiments, respectively.

## Author Contributions

Conceptualization: S.S., V.M., D.A.; Methodology: S.S., S.B., V.M., D.A.; Software: S.S., M.C.T., V.M.; Validation: S.S., S.B.; Formal analysis: S.S., S.B., U.A, M.C.T., M.D.G., A.K., C.R.W., G.P., M.Z.P.; Investigation: S.S., S.B., U.A, M.C.T., M.D.G., A.K., C.R.W., G.P., M.Z.P.; Resources: V.M., R.A.B. D.A.; Data curation: S.S., S.B., V.M., D.A.; Writing – revised draft: S.S., S.B., V.M., D.A.; Writing & editing: S.S., S.B., M.C.T., V.M., D.A.; Visualization: S.S., S.B., C.W.; Supervision: V.M., D.A.; Project administration: D.A.; Funding acquisition: V.M., D.A.

## MATERIALS AND CORRESPONDENCE

The materials are available upon request to Dritan Agalliu.

## REFERENCES

1 Daneman, R. & Engelhardt, B. Brain barriers in health and disease. Neurobiology of disease 107, 1–3 (2017).

2 Liebner, S. et al. Functional morphology of the blood-brain barrier in health and disease. Acta Neuropathol 135, 311–336, doi:10.1007/s00401-018-1815-1 (2018).

3 Kirk, J., Plumb, J., Mirakhur, M. & McQuaid, S. Tight junctional abnormality in multiple sclerosis white matter affects all calibres of vessel and is associated with blood-brain barrier leakage and active demyelination. The Journal of pathology 201, 319–327 (2003).

4 Pan, W., Banks, W. A., Kennedy, M. K., Gutierrez, E. G. & Kastin, A. J. Differential permeability of the BBB in acute EAE: enhanced transport of TNT-alpha. The American journal of physiology 271, E636–642 (1996).

5 Brown, A., McFarlin, D. E. & Raine, C. S. Chronologic neuropathology of relapsing experimental allergic encephalomyelitis in the mouse. Laboratory investigation; a journal of technical methods and pathology 46, 171–185 (1982).

6 Alvarez, J. I. et al. Focal disturbances in the blood-brain barrier are associated with formation of neuroinflammatory lesions. Neurobiol Dis 74, 14–24, doi:10.1016/j.nbd.2014.09.016 (2015).

7 Lossinsky, A. S., Badmajew, V., Robson, J. A., Moretz, R. C. & Wisniewski, H. M. Sites of egress of inflammatory cells and horseradish peroxidase transport across the blood-brain barrier in a murine model of chronic relapsing experimental allergic encephalomyelitis. Acta Neuropathol 78, 359–371 (1989).

8 Maggi, P. et al. The formation of inflammatory demyelinated lesions in cerebral white matter. Ann Neurol 76, 594–608, doi:10.1002/ana.24242 (2014).

9 Lutz, S. E. et al. Caveolin1 is required for Th1 cell infiltration, but not tight junction remodeling, at the blood-brain barrier in autoimmune neuroinflammation. Cell reports 21, 2104–2117 (2017).

10 Plumb, J., McQuaid, S., Mirakhur, M. & Kirk, J. Abnormal endothelial tight junctions in active lesions and normal-appearing white matter in multiple sclerosis. *Brain pathology (Zurich*, Switzerland) 12, 154–169 (2002).

11 Kermode, A. G. et al. Breakdown of the blood-brain barrier precedes symptoms and other MRI signs of new lesions in multiple sclerosis. Pathogenetic and clinical implications. Brain : a journal of neurology 113 (Pt 5), 1477–1489 (1990).

12 Reijerkerk, A. et al. Tissue-type plasminogen activator is a regulator of monocyte diapedesis through the brain endothelial barrier. Journal of immunology (Baltimore, Md. :1950) 181, 3567–3574 (2008).

13 Winger, R. C., Koblinski, J. E., Kanda, T., Ransohoff, R. M. & Muller, W. A. Rapid remodeling of tight junctions during paracellular diapedesis in a human model of the blood-brain barrier. Journal of immunology (Baltimore, Md. : 1950) 193, 2427–2437, doi:10.4049/jimmunol.1400700 (2014).

14 Aube, B. et al. Neutrophils mediate blood-spinal cord barrier disruption in demyelinating neuroinflammatory diseases. Journal of immunology (Baltimore, Md. : 1950) 193, 2438–2454, doi:10.4049/jimmunol.1400401 (2014).

15 Abadier, M. et al. Cell surface levels of endothelial ICAM-1 influence the transcellular or paracellular T-cell diapedesis across the blood-brain barrier. European journal of immunology 45, 1043–1058, doi:10.1002/eji.201445125 (2015).

16 Lake, B. B. et al. Integrative single-cell analysis of transcriptional and epigenetic states in the human adult brain. Nature Biotechnology 36, 70–80, doi:10.1038/nbt.4038 (2018).

17 Biswas, S., Cottarelli, A. & Agalliu, D. Neuronal and glial regulation of CNS angiogenesis and barriergenesis. Development 147, dev182279 (2020).

18 Girolamo, F., Coppola, C., Ribatti, D. & Trojano, M. Angiogenesis in multiple sclerosis and experimental autoimmune encephalomyelitis. Acta neuropathologica communications 2, 1–17 (2014).

19 Lengfeld, J., Cutforth, T. & Agalliu, D. The role of angiogenesis in the pathology of multiple sclerosis. Vascular cell 6, 1–6 (2014).

20 Proescholdt, M. A., Jacobson, S., Tresser, N., Oldfield, E. H. & Merrill, M. J. Vascular endothelial growth factor is expressed in multiple sclerosis plaques and can induce inflammatory lesions in experimental allergic encephalomyelitis rats. Journal of neuropathology and experimental neurology 61, 914–925 (2002).

21 Holley, J. E., Newcombe, J., Whatmore, J. L. & Gutowski, N. J. Increased blood vessel density and endothelial cell proliferation in multiple sclerosis cerebral white matter. Neurosci Lett 470, 65–70, doi:10.1016/j.neulet.2009.12.059 (2010).

22 Seabrook, T. J. et al. Angiogenesis is present in experimental autoimmune encephalomyelitis and pro-angiogenic factors are increased in multiple sclerosis lesions. J Neuroinflammation 7, 95, doi:10.1186/1742-2094-7-95 (2010).

23 Kirk, S. L. & Karlik, S. J. VEGF and vascular changes in chronic neuroinflammation. J Autoimmun 21, 353–363 (2003).

24 Roscoe, W. A., Welsh, M. E., Carter, D. E. & Karlik, S. J. VEGF and angiogenesis in acute and chronic MOG((35-55)) peptide induced EAE. J Neuroimmunol 209, 6–15, doi:10.1016/j.jneuroim.2009.01.009 (2009).

25 Boroujerdi, A., Welser-Alves, J. V. & Milner, R. Extensive vascular remodeling in the spinal cord of pre-symptomatic experimental autoimmune encephalomyelitis mice; increased vessel expression of fibronectin and the alpha5beta1 integrin. Exp Neurol 250, 43–51, doi:10.1016/j.expneurol.2013.09.009 (2013).

26 MacMillan, C. J. et al. Bevacizumab diminishes experimental autoimmune encephalomyelitis by inhibiting spinal cord angiogenesis and reducing peripheral T-cell responses. J Neuropathol Exp Neurol 71, 983–999, doi:10.1097/NEN.0b013e3182724831 (2012).

27 Buch, S. et al. Revealing vascular abnormalities and measuring small vessel density in multiple sclerosis lesions using USPIO. Neuroimage Clin 29, 102525, doi:10.1016/j.nicl.2020.102525 (2021).

28 MacMillan, C. J. et al. Murine experimental autoimmune encephalomyelitis is diminished by treatment with the angiogenesis inhibitors B20-4.1. 1 and angiostatin (K1-3). PLoS One 9, e89770 (2014).

29 Matsumoto, K. & Ema, M. Roles of VEGF-A signalling in development, regeneration, and tumours. The Journal of Biochemistry 156, 1–10 (2014).

30 Carmeliet, P. VEGF as a key mediator of angiogenesis in cancer. Oncology 69, 4–10 (2005).

31 Campochiaro, P. A. Molecular pathogenesis of retinal and choroidal vascular diseases. Progress in retinal and eye research 49, 67–81 (2015).

32 Su, J. J. et al. Upregulation of vascular growth factors in multiple sclerosis: correlation with MRI findings. Journal of the neurological sciences 243, 21–30 (2006).

33 Presta, L. G. et al. Humanization of an anti-vascular endothelial growth factor monoclonal antibody for the therapy of solid tumors and other disorders. Cancer research 57, 4593–4599 (1997).

34 Ferrara, N., Hillan, K. J., Gerber, H.-P. & Novotny, W. Discovery and development of bevacizumab, an anti-VEGF antibody for treating cancer. Nature reviews Drug discovery 3, 391–400 (2004).

35 Ferrara, N., Hillan, K. J. & Novotny, W. Bevacizumab (Avastin), a humanized anti-VEGF monoclonal antibody for cancer therapy. Biochemical and biophysical research communications 333, 328–335 (2005).

36 Li, Y.-L., Zhao, H. & Ren, X.-B. Relationship of VEGF/VEGFR with immune and cancer cells: staggering or forward? Cancer biology & medicine 13, 206 (2016).

37 Daneman, R. et al. Wnt/β-catenin signaling is required for CNS, but not non-CNS, angiogenesis. Proceedings of the National Academy of Sciences 106, 641–646 (2009).

38 Stenman, J. M. et al. Canonical Wnt signaling regulates organ-specific assembly and differentiation of CNS vasculature. Science 322, 1247–1250 (2008).

39 Liebner, S. et al. Wnt/β-catenin signaling controls development of the blood–brain barrier. Journal of Cell Biology 183, 409–417 (2008).

40 Zhou, Y. et al. Canonical WNT signaling components in vascular development and barrier formation. The Journal of clinical investigation 124, 3825–3846 (2014).

41 Tran, K. A. et al. Endothelial β-catenin signaling is required for maintaining adult blood– brain barrier integrity and central nervous system homeostasis. Circulation 133, 177–186 (2016).

42 Lengfeld, J. E. et al. Endothelial Wnt/beta-catenin signaling reduces immune cell infiltration in multiple sclerosis. Proceedings of the National Academy of Sciences of the United States of America, doi:10.1073/pnas.1609905114 (2017).

43 Sullivan, L. A. et al. r84, a novel therapeutic antibody against mouse and human VEGF with potent anti-tumor activity and limited toxicity induction. PloS one 5, e12031 (2010).

44 Roland, C. L. et al. Inhibition of vascular endothelial growth factor reduces angiogenesis and modulates immune cell infiltration of orthotopic breast cancer xenografts. Mol Cancer Ther 8, 1761–1771, doi:10.1158/1535-7163.MCT-09-0280 (2009).

45 Hao, Y. et al. Integrated analysis of multimodal single-cell data. Cell (2021).

46 Vanlandewijck, M. et al. A molecular atlas of cell types and zonation in the brain vasculature. Nature 554, 475–480 (2018).

47 Miao, Z. et al. Putative cell type discovery from single-cell gene expression data. Nature methods 17, 621–628, doi:10.1038/s41592-020-0825-9 (2020).

48 Halpern, K. B. et al. Single-cell spatial reconstruction reveals global division of labour in the mammalian liver. Nature 542, 352–356 (2017).

49 Korsunsky, I. et al. Fast, sensitive and accurate integration of single-cell data with Harmony. Nature methods 16, 1289–1296, doi:10.1038/s41592-019-0619-0 (2019).

50 Munji, R. N. et al. Profiling the mouse brain endothelial transcriptome in health and disease models reveals a core blood-brain barrier dysfunction module. Nat Neurosci 22, 1892–1902, doi:10.1038/s41593-019-0497-x (2019).

51 Jeong, H.-W. et al. Single-cell transcriptomics reveals functionally specialized vascular endothelium in brain. eLife 11, e57520, doi:10.7554/eLife.57520 (2022).

52 Subramanian, A. et al. Gene set enrichment analysis: a knowledge-based approach for interpreting genome-wide expression profiles. Proceedings of the National Academy of Sciences 102, 15545–15550 (2005).

53 Zarkada, G. et al. Specialized endothelial tip cells guide neuroretina vascularization and blood-retina-barrier formation. Developmental Cell 56, 2237–2251. e2236 (2021).

54 Zhao, Q. et al. Single-cell transcriptome analyses reveal endothelial cell heterogeneity in tumors and changes following antiangiogenic treatment. Cancer research 78, 2370–2382 (2018).

55 Conway, E. M., Collen, D. & Carmeliet, P. Molecular mechanisms of blood vessel growth. Cardiovascular research 49, 507–521 (2001).

56 Potenta, S., Zeisberg, E. & Kalluri, R. The role of endothelial-to-mesenchymal transition in cancer progression. British journal of cancer 99, 1375–1379 (2008).

57 Platel, V., Faure, S., Corre, I. & Clere, N. Endothelial-to-mesenchymal transition (EndoMT): roles in tumorigenesis, metastatic extravasation and therapy resistance. Journal of oncology 2019 (2019).

58 Neve, A., Cantatore, F. P., Maruotti, N., Corrado, A. & Ribatti, D. Extracellular matrix modulates angiogenesis in physiological and pathological conditions. BioMed research international 2014 (2014).

59 Clere, N., Renault, S. & Corre, I. Endothelial-to-mesenchymal transition in cancer. Frontiers in Cell and Developmental Biology 8 (2020).

60 Hogan, K. A., Ambler, C. A., Chapman, D. L. & Bautch, V. L. The neural tube patterns vessels developmentally using the VEGF signaling pathway. (2004).

61 James, J. M., Gewolb, C. & Bautch, V. L. Neurovascular development uses VEGF-A signaling to regulate blood vessel ingression into the neural tube. (2009).

62 Nguyen, H.-L. et al. TGF-β signaling in endothelial cells, but not neuroepithelial cells, is essential for cerebral vascular development. Laboratory investigation 91, 1554–1563 (2011).

63 Goveia, J. et al. An integrated gene expression landscape profiling approach to identify lung tumor endothelial cell heterogeneity and angiogenic candidates. Cancer cell 37, 21–36. e13 (2020).

64 Hupe, M. et al. Gene expression profiles of brain endothelial cells during embryonic development at bulk and single-cell levels. Science Signaling 10 (2017).

65 Sabbagh, M. F. et al. Transcriptional and epigenomic landscapes of CNS and non-CNS vascular endothelial cells. elife 7, e36187 (2018).

66 del Toro, R. et al. Identification and functional analysis of endothelial tip cell–enriched genes. *Blood*, The Journal of the American Society of Hematology 116, 4025–4033 (2010).

67 Wang, H. U., Chen, Z.-F. & Anderson, D. J. Molecular distinction and angiogenic interaction between embryonic arteries and veins revealed by ephrin-B2 and its receptor Eph-B4. Cell 93, 741–753 (1998).

68 Davies, M. H., Stempel, A. J., Hubert, K. E. & Powers, M. R. Altered vascular expression of EphrinB2 and EphB4 in a model of oxygen-induced retinopathy. Developmental Dynamics 239, 1695–1707 (2010).

69 Crist, A. M., Young, C. & Meadows, S. M. Characterization of arteriovenous identity in the developing neonate mouse retina. Gene Expression Patterns 23, 22–31 (2017).

70 Zhang, G., Yang, X. & Gao, R. Research progress on the structure and function of endomucin. Animal Models and Experimental Medicine 3, 325–329 (2020).

71 Park-Windhol, C. et al. Endomucin inhibits VEGF-induced endothelial cell migration, growth, and morphogenesis by modulating VEGFR2 signaling. Scientific reports 7, 1–13 (2017).

72 Coelho-Santos, V., Berthiaume, A.-A., Ornelas, S., Stuhlmann, H. & Shih, A. Y. Imaging the construction of capillary networks in the neonatal mouse brain. Proceedings of the National Academy of Sciences 118 (2021).

73 Uemura, A., Kusuhara, S., Katsuta, H. & Nishikawa, S.-I. Angiogenesis in the mouse retina: a model system for experimental manipulation. Experimental cell research 312, 676–683 (2006).

74 Lee, H.-W. et al. The Role of Venous Endothelial Cells in Developmental and Pathologic Angiogenesis. Circulation (2021).

75 Schmidt, M., et al. EGFL7 regulates the collective migration of endothelial cells by restricting their spatial distribution. (2007).

76 Parker, L. H. et al. The endothelial-cell-derived secreted factor Egfl7 regulates vascular tube formation. Nature 428, 754–758 (2004).

77 Nichol, D. et al. Impaired angiogenesis and altered Notch signaling in mice overexpressing endothelial Egfl7. *Blood*, The Journal of the American Society of Hematology 116, 6133–6143 (2010).

78 Poissonnier, L., Villain, G., Soncin, F. & Mattot, V. Egfl7 is differentially expressed in arteries and veins during retinal vascular development. PloS one 9, e90455 (2014).

79 Joshkon, A. et al. Role of CD146 (MCAM) in Physiological and Pathological Angiogenesis—Contribution of New Antibodies for Therapy. Biomedicines 8, 633 (2020).

80 Jiang, T. et al. CD146 is a coreceptor for VEGFR-2 in tumor angiogenesis. *Blood*, The Journal of the American Society of Hematology 120, 2330–2339 (2012).

81 Lee, K.-m., et al. Extracellular matrix protein 1 regulates cell proliferation and trastuzumab resistance through activation of epidermal growth factor signaling. Breast Cancer Research 16, 1–17 (2014).

82 De Luca, A. et al. The role of the EGFR signaling in tumor microenvironment. Journal of cellular physiology 214, 559–567 (2008).

83 Stoppelli, M. P. in Madame Curie Bioscience Database [Internet] (Landes Bioscience, 2013).

84 Takayama, Y. et al. Inhibition of PAI-1 limits tumor angiogenesis regardless of angiogenic stimuli in malignant pleural mesothelioma. Cancer research 76, 3285–3294 (2016).

85 Girolamo, F., Coppola, C., Ribatti, D. & Trojano, M. Angiogenesis in multiple sclerosis and experimental autoimmune encephalomyelitis. Acta Neuropathol Commun 2, 84, doi:10.1186/s40478-014-0084-z (2014).

86 Kaufmann, M. et al. Identification of early neurodegenerative pathways in progressive multiple sclerosis. Nat Neurosci 25, 944–955, doi:10.1038/s41593-022-01097-3 (2022).

87 Kim, B. R. et al. MARCKSL1 exhibits anti-angiogenic effects through suppression of VEGFR-2-dependent Akt/PDK-1/mTOR phosphorylation. Oncol Rep 35, 1041–1048, doi:10.3892/or.2015.4408 (2016).

88 Venkatraman, L. & Claesson-Welsh, L. in Tumor Angiogenesis: A Key Target for Cancer Therapy (ed Dieter Marmé) 33–50 (Springer International Publishing, 2019).

89 Potente, M., Gerhardt, H. & Carmeliet, P. Basic and therapeutic aspects of angiogenesis. Cell 146, 873–887, doi:10.1016/j.cell.2011.08.039 (2011).

90 Potente, M. & Carmeliet, P. The Link Between Angiogenesis and Endothelial Metabolism. Annu Rev Physiol 79, 43–66, doi:10.1146/annurev-physiol-021115-105134 (2017).

91 Rohlenova, K. et al. Single-Cell RNA Sequencing Maps Endothelial Metabolic Plasticity in Pathological Angiogenesis. Cell Metab 31, 862–877 e814, doi:10.1016/j.cmet.2020.03.009 (2020).

92 Gavalas, N. G. et al. VEGF directly suppresses activation of T cells from ascites secondary to ovarian cancer via VEGF receptor type 2. British Journal of Cancer 107, 1869–1875, doi:10.1038/bjc.2012.468 (2012).

93 Bourhis, M., Palle, J., Galy-Fauroux, I. & Terme, M. Direct and Indirect Modulation of T Cells by VEGF-A Counteracted by Anti-Angiogenic Treatment. Front Immunol 12, 616837, doi:10.3389/fimmu.2021.616837 (2021).

94 Jetten, N. et al. Anti-inflammatory M2, but not pro-inflammatory M1 macrophages promote angiogenesis in vivo. Angiogenesis 17, 109–118 (2014).

95 Quiding-Järbrink, M., Smith, D. A. & Bancroft, G. J. Production of matrix metalloproteinases in response to mycobacterial infection. Infection and immunity 69, 5661–5670, doi:10.1128/iai.69.9.5661-5670.2001 (2001).

96 Conway, D. E. & Schwartz, M. A. Mechanotransduction of shear stress occurs through changes in VE-cadherin and PECAM-1 tension: implications for cell migration. Cell Adh Migr 9, 335–339, doi:10.4161/19336918.2014.968498 (2015).

97 Argaw, A. T., Gurfein, B. T., Zhang, Y., Zameer, A. & John, G. R. VEGF-mediated disruption of endothelial CLN-5 promotes blood-brain barrier breakdown. Proceedings of the National Academy of Sciences 106, 1977–1982 (2009).

98 Antonetti, D. A., Barber, A. J., Hollinger, L. A., Wolpert, E. B. & Gardner, T. W. Vascular endothelial growth factor induces rapid phosphorylation of tight junction proteins occludin and zonula occluden 1: a potential mechanism for vascular permeability in diabetic retinopathy and tumors. Journal of Biological Chemistry 274, 23463–23467 (1999).

99 Murakami, T., Felinski, E. A. & Antonetti, D. A. Occludin phosphorylation and ubiquitination regulate tight junction trafficking and vascular endothelial growth factor-induced permeability. Journal of Biological Chemistry 284, 21036–21046 (2009).

100 Wang, W., Dentler, W. L. & Borchardt, R. T. VEGF increases BMEC monolayer permeability by affecting occludin expression and tight junction assembly. American Journal of Physiology-Heart and Circulatory Physiology 280, H434–H440 (2001).

101 Gavard, J. & Gutkind, J. S. VEGF controls endothelial-cell permeability by promoting the β-arrestin-dependent endocytosis of VE-cadherin. Nature cell biology 8, 1223–1234 (2006).

102 Senger, D. R. et al. Tumor cells secrete a vascular permeability factor that promotes accumulation of ascites fluid. Science 219, 983–985 (1983).

103 Lopes Pinheiro, M. A., et al. Immune cell trafficking across the barriers of the central nervous system in multiple sclerosis and stroke. Biochimica et biophysica acta, doi:10.1016/j.bbadis.2015.10.018 (2015).

104 Argaw, A. T. et al. Astrocyte-derived VEGF-A drives blood-brain barrier disruption in CNS inflammatory disease. J Clin Invest 122, 2454–2468, doi:10.1172/jci60842 (2012).

105 Vanlandewijck, M., Lebouvier, T., Mäe, M. A., Nahar, K. & Betsholtz, C. Primary isolation of vascular cells from murine brain for single cell sequencing. (2018).

106 Yuan, J. & Sims, P. A. An Automated Microwell Platform for Large-Scale Single Cell RNA-Seq. Sci Rep 6, 33883, doi:10.1038/srep33883 (2016).

107 Yuan, J. et al. Single-cell transcriptome analysis of lineage diversity in high-grade glioma. Genome Med 10, 57, doi:10.1186/s13073-018-0567-9 (2018).

108 Menon, V. Clustering single cells: a review of approaches on high-and low-depth single-cell RNA-seq data. Brief Funct Genomics 17, 240–245, doi:10.1093/bfgp/elx044 (2018).

109 Tasic, B. et al. Adult mouse cortical cell taxonomy revealed by single cell transcriptomics. Nat Neurosci 19, 335–346, doi:10.1038/nn.4216 (2016).

110 Sorensen, S. A. et al. Correlated gene expression and target specificity demonstrate excitatory projection neuron diversity. Cereb Cortex 25, 433–449, doi:10.1093/cercor/bht243 (2015).

111 Hawrylycz, M. et al. Canonical genetic signatures of the adult human brain. Nat Neurosci 18, 1832–1844, doi:10.1038/nn.4171 (2015).

112 Mazzoni, J. et al. The Wnt Inhibitor Apcdd1 Coordinates Vascular Remodeling and Barrier Maturation of Retinal Blood Vessels. Neuron 96, 1055–1069 e1056, doi:10.1016/j.neuron.2017.10.025 (2017).

