## Extended Data Figures and Legends for "Single-cell RNA-sequencing implicates venous endothelial cells as a source of VEGF-A-mediated neo-angiogenesis in neuroinflammation"

*equal contribution first authors

**equal contribution senior authors

Mailing address:

Columbia University Irving Medical Center

650 West 168th Street

Black Building Room 310

New York, NY 10032

ORCID: 0000-0002-5375-4143

### Supporting Information - Inventory

#### I. Extended Data Figures and Figure Legends

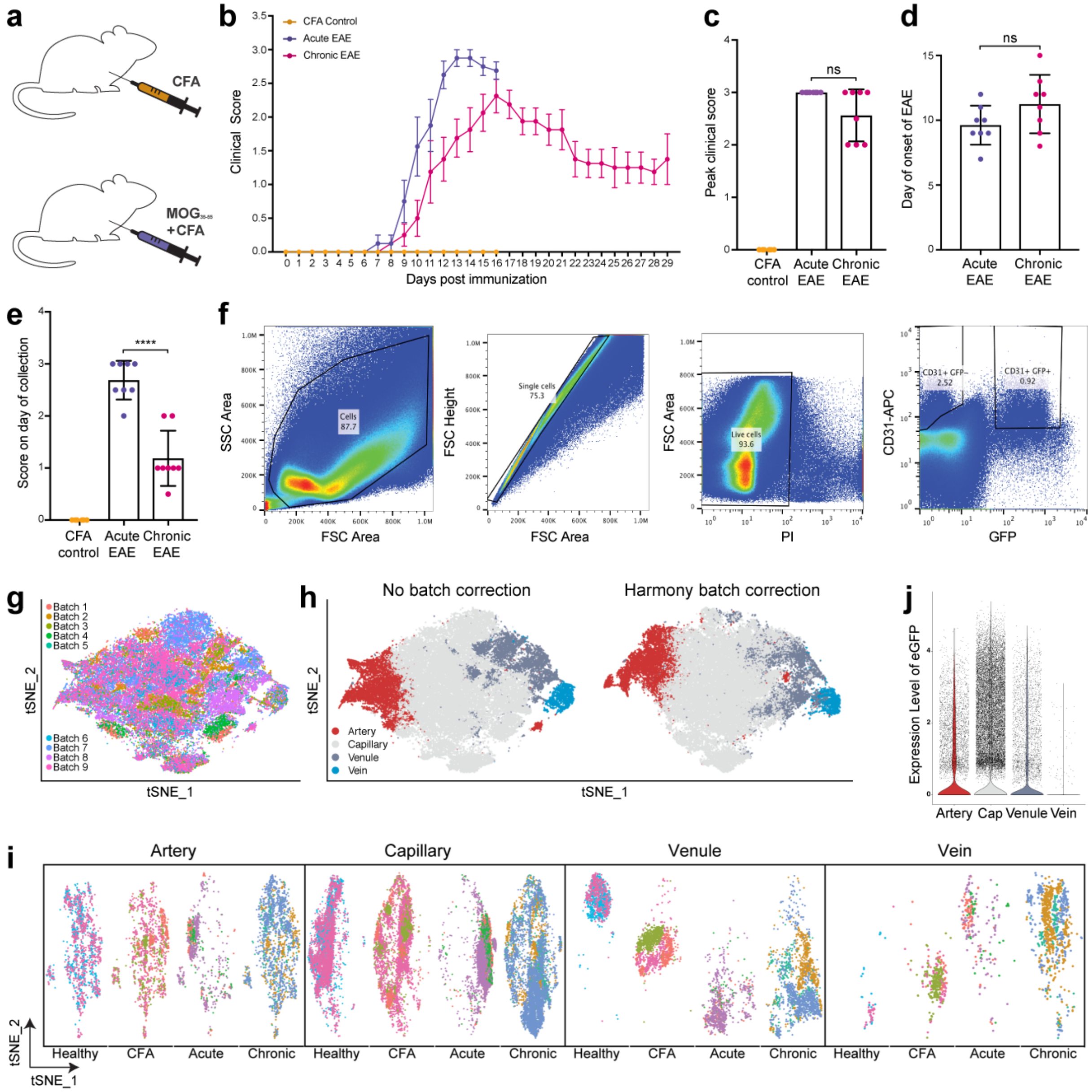


**Extended Data Figure 1. Disease course of MOG35-55 EAE mice used for single-cell RNA-sequencing, gating strategy for EC isolation and data analysis. a**) Mice were immunized with either MOG35-55 peptide and CFA/Ptx to induce EAE, or CFA/Ptx (CFA controls). **b-e**) Clinical EAE score curves (b), peak clinical scores (c), day of disease onset (d), and score on the day of collection (e) for CFA control (yellow), acute (16 d.p.i., purple) and chronic (29d.p.i., magenta)

EAE cohorts used for single-cell RNA-sequencing (scRNA-seq). Two mice were pooled for each sample to obtain sufficient endothelial cells (ECs) for scRNA-seq. Data represent mean ± SEM; n = 6 female CFA/PBS control (3 samples), n = 8 female acute EAE (4 samples), n = 8 female chronic EAE mice (4 samples); * $p<0.05$, ** $p<0.01$; **** $p<0.0001$; b) repeated measure ANOVA with Bonferroni correction for multiple testing; c, e) one-way ANOVA with Bonferroni correction for multiple testing; d) Student's unpaired two-sided t-test. **f**) FACS plots show the gating strategy for sorting spinal cord ECs from either control [healthy or CFA; n = 6 mice (3 samples per condition)], acute or chronic *Tcf/Lef*::H2B::eGFP (Wnt reporter) EAE mice [n = 8 mice (4 samples per condition)] for scRNA-seq. $eGFP^+$ and $eGFP^-$ spinal cord ECs were pooled prior to performing scRNA-seq from all conditions. **g**) t-distributed Stochastic Neighbor Embedding (t-SNE) plot, color-coded for each batch. **h**) t-SNE plot, color coded for each EC subtype, with and without Harmony batch correction. Re-embedding into t-SNE space after applying Harmony correction does not show any dramatic differences visually in the overall organization of EC clusters. **i**) Individually re-clustered t-SNE plots for each EC subtype (artery, capillary, venule, vein), color-coded by batch, grouped by disease state (healthy, CFA control, acute and chronic EAE). These plots demonstrate that all ECs subtypes from each distinct disease state cluster together irrespective of the batch, suggesting presence of minimal batch effects in the dataset. **j**) Violin plot shows very low expression of *eGFP* mRNA in vECs isolated from *Tcf/Lef*::H2B::eGFP EAE mice, indicative of low-to-absent Wnt/β-catenin signaling activity in vein ECs during EAE.

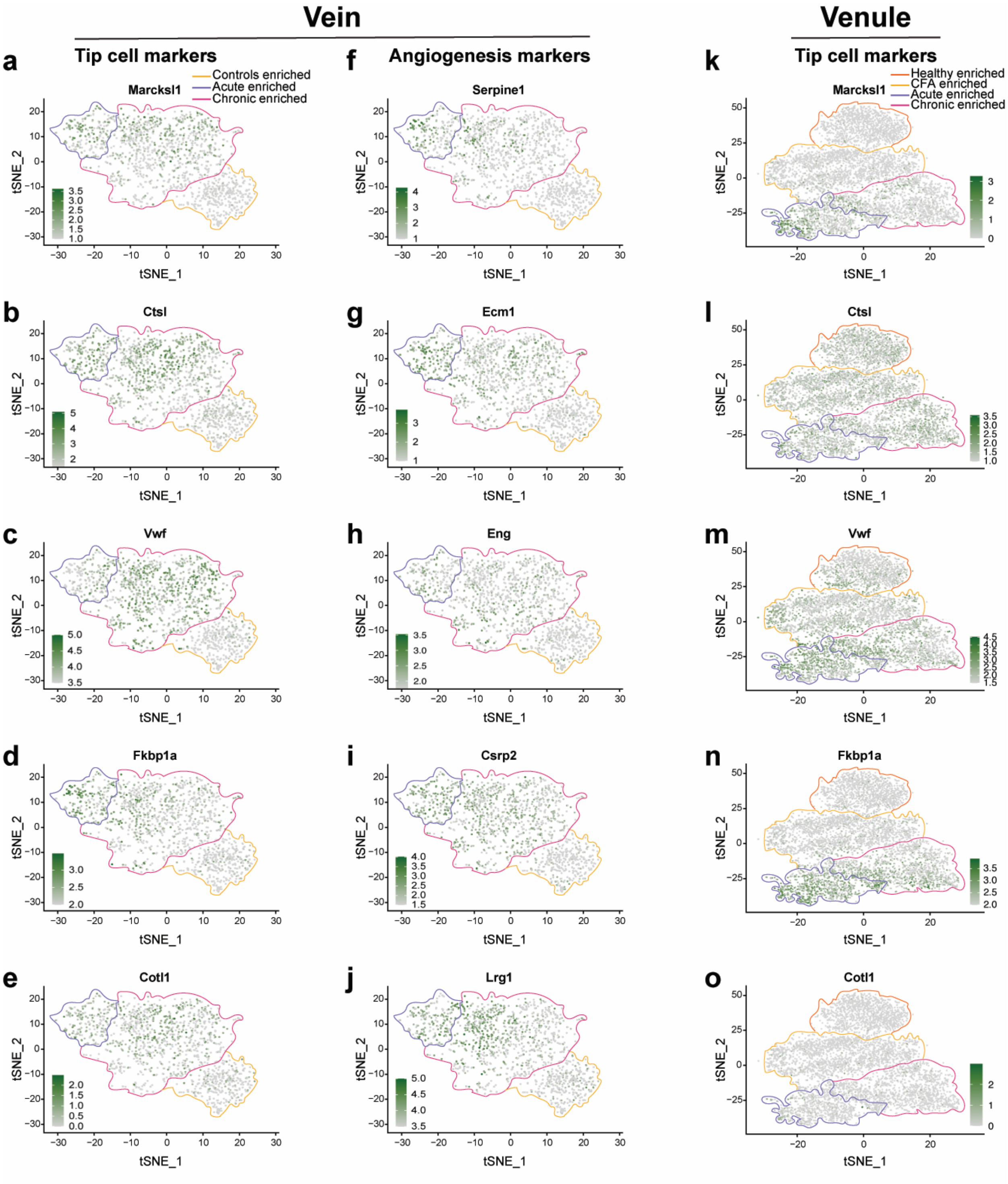


**Extended Data Figure 2. Angiogenic and tip cell marker transcripts are upregulated in vECs and vnECs in both acute and chronic EAE. a-j**) Feature plots show expression of angiogenic and tip cell markers in spinal cord vein ECs. The expression of angiogenic and tip cell markers is increased in vECs isolated from both acute and chronic EAE conditions. **k-o**) Feature plots show expression of tip cell markers in spinal cord venule ECs. The expression of tip cell markers is increased in vnECs isolated from both acute and chronic EAE conditions.

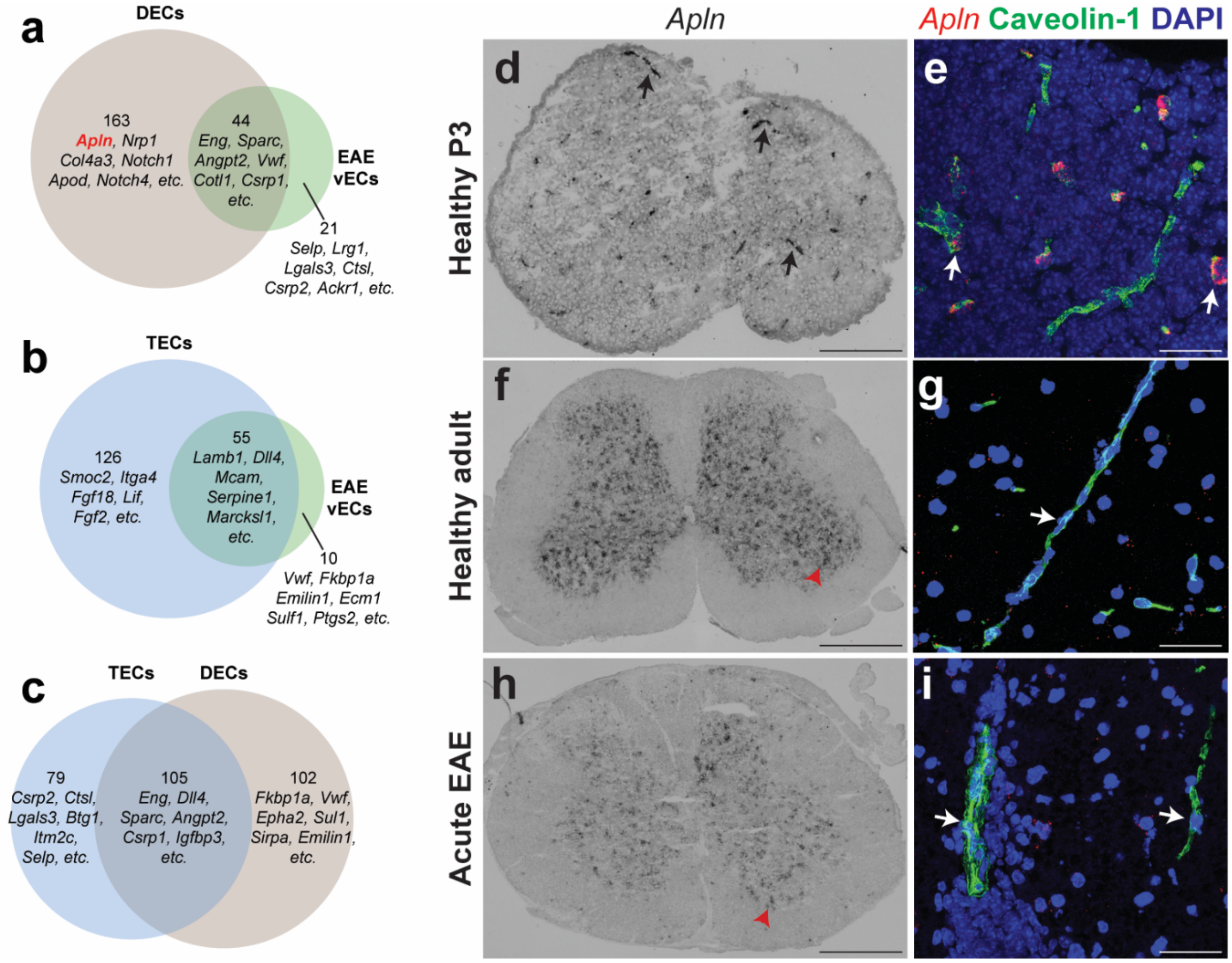


**Extended Data Figure 3. Congruency in angiogenic gene expression among acute EAE vECs, developing CNS endothelial cells (DECs), and lung tumor endothelial cells (TECs). a-c**) Venn diagrams of the overlap in angiogenic gene signatures between DECs and EAE vECs (a), TECs and EAE vECs (b), and TECs and DECs (c). Venn diagrams show the numbers of common and unique genes between the two populations in each comparison, as well as some of the key gene names. **d, f, h**) Alkaline Phosphatase *in situ* hybridization for *Apln* mRNA, and **e, g, i**) fluorescent *in situ* hybridization for *Apln* mRNA combined with immunofluorescence for Caveolin-1 (EC marker) and DAPI, in either developing (P3; d, e), adult (f, g) healthy spinal cords, and acute EAE spinal cords (16 d.p.i.; score 2.5; h, g). *Apln* mRNA is expressed by many blood vessels in the developing P3 spinal cord (d, black arrows; e, white arrows) when CNS angiogenesis is ongoing, but is absent in the adult spinal cord (g, white arrow) when CNS angiogenesis is completed. *Apln* mRNA is expressed in neurons in the adult spinal cord (f, h, shown with a red arrowhead). *Alpn* mRNA expression is absent from blood vessels located in white matter lesions in acute EAE spinal cords (i), although other angiogenesis markers are expressed in these vessels (see Figure 5). Scale bars: a = 260 µm; c,e = 575 µm; b,d,f = 40 µm.

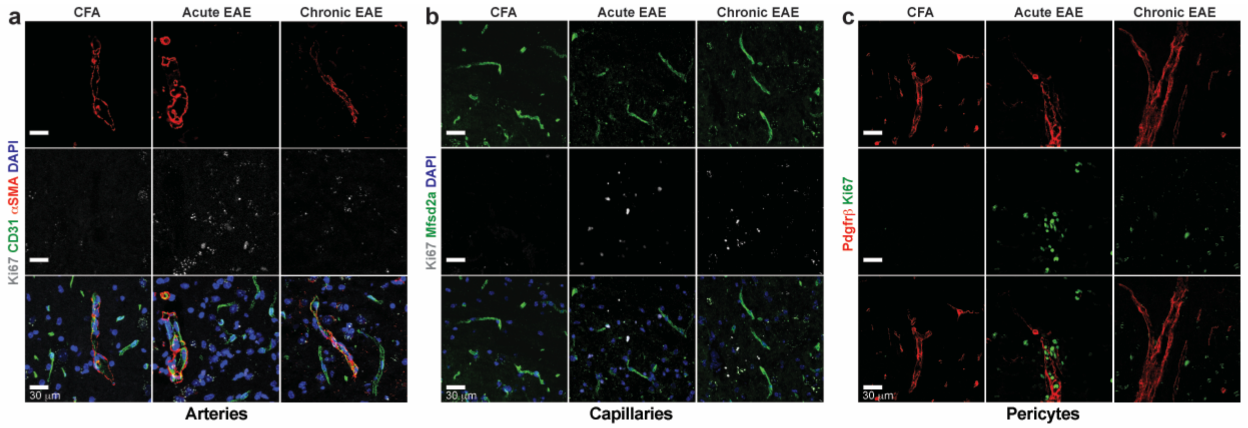


**Extended Data Figure 4. Arterial ECs, capillary ECs and pericytes do not proliferate in EAE.** **a)** Sagittal sections of control (CFA), acute (16 d.p.i) and chronic EAE (28 d.p.i) spinal cords were immunostained with antibodies against alpha-smooth muscle actin (αSMA) to labels arteries, Ki67 (proliferation marker), CD31 (EC marker) and DAPI. There are no proliferative arterial ECs in EAE. **b)** Sagittal sections of control (CFA), acute and chronic EAE spinal cords were immunostained with antibodies against Mfsd2a (capillary ECs), Ki67 and DAPI. There are no proliferative capillary ECs in EAE. c) Sagittal sections of control (CFA), acute and chronic EAE spinal cords were immunostained for Pdgfrβ (pericyte marker) and Ki67. Pericytes do not proliferate in EAE. Scale bars = 30 µm.

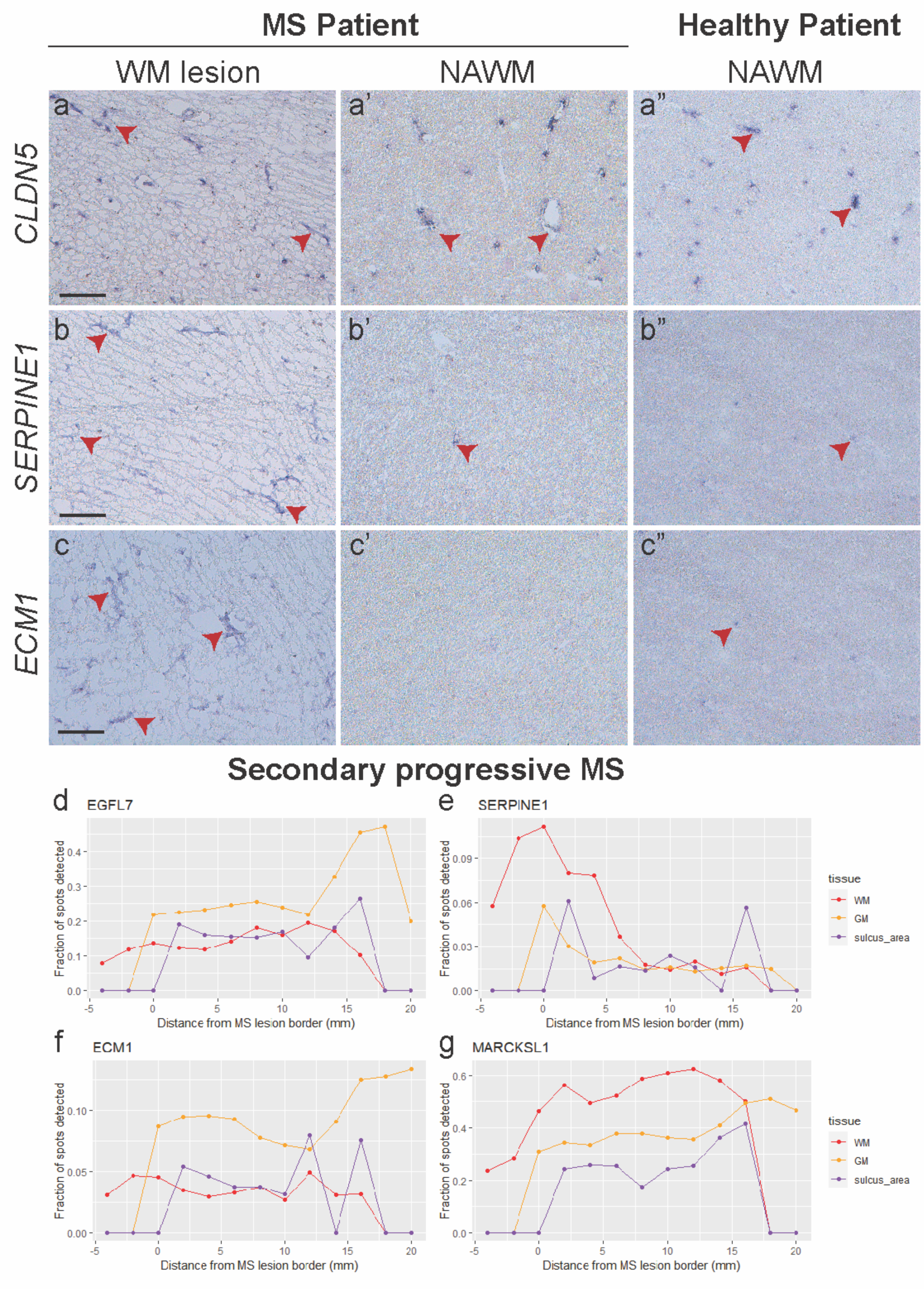

MS Patient
Healthy Patient
WM lesion
NAWM
NAWM
CLDN5
SERPINE1
ECM1
a
a'
a"
b
b'
b"
c
c'
c"
Secondary progressive MS
d
EGFL7
e
SERPINE1
f
ECM1
g
MARCKSL1
Fraction of spots detected
Distance from MS lesion border (mm)
tissue
WM
GM
sulcus_area

**Extended Data Figure 5. Increased expression of angiogenic markers in human relapsing remitting and secondary progressive multiple sclerosis cases. a-c”**) *In situ* hybridization for human *CLAUDIN-5* (a-a”), *SERPINE1* (b-b”), and *ECM1* (c-c”) mRNAs show expression of the angiogenesis markers *SERPINE1* and *ECM1* mRNAs in white matter (WM) multiple sclerosis (MS) lesions, but neither in the normal appearing white matter (NAWM) from MS patients, nor in the NAWM from non-neurological control cases. *CLAUDIN-5* mRNA is expressed by all blood vessels in human brain tissues. The fresh frozen MS brain samples were obtained from relapsing-remitting MS (RRMS) cases (N=4), and non-neurological controls (N=4). Scale bars = 40 µm. **d-g**) Line plots summarizing the detection of four angiogenesis mRNAs for *EGFL7* (d), *SERPINE1* (e), *ECM1* (f) and *MARCKSL1* (g) in either white matter (WM; red line), gray matter (GM, orange), or the sulcus area (purple line), with respect to the distance from the MS lesion border in secondary progressive MS cases. All expression data were obtained from Kaufmann et al., 2022[1], which used the original spatial transcriptomics platform to profile gene expression within an array of 100-µm diameter spots from MS lesion borders. Methods to measure distance from lesion borders are outlined in the publication[1], and were imported in the same way for this analysis. Here, we binned all spots by their distance from MS lesion border in each tissue type, using bins of 0.5 mm width, and calculated the fraction of spots within each bin where the four angiogenesis genes were detected. White matter spots far away from MS lesions show lower detection for these genes, with *SERPINE1* and *MARCKSL1* mRNAs showing the most marked effect of all four angiogenesis genes. These mRNAs are expressed at very high levels near the border lesion in the white matter, but not either gray matter or sulcus region.

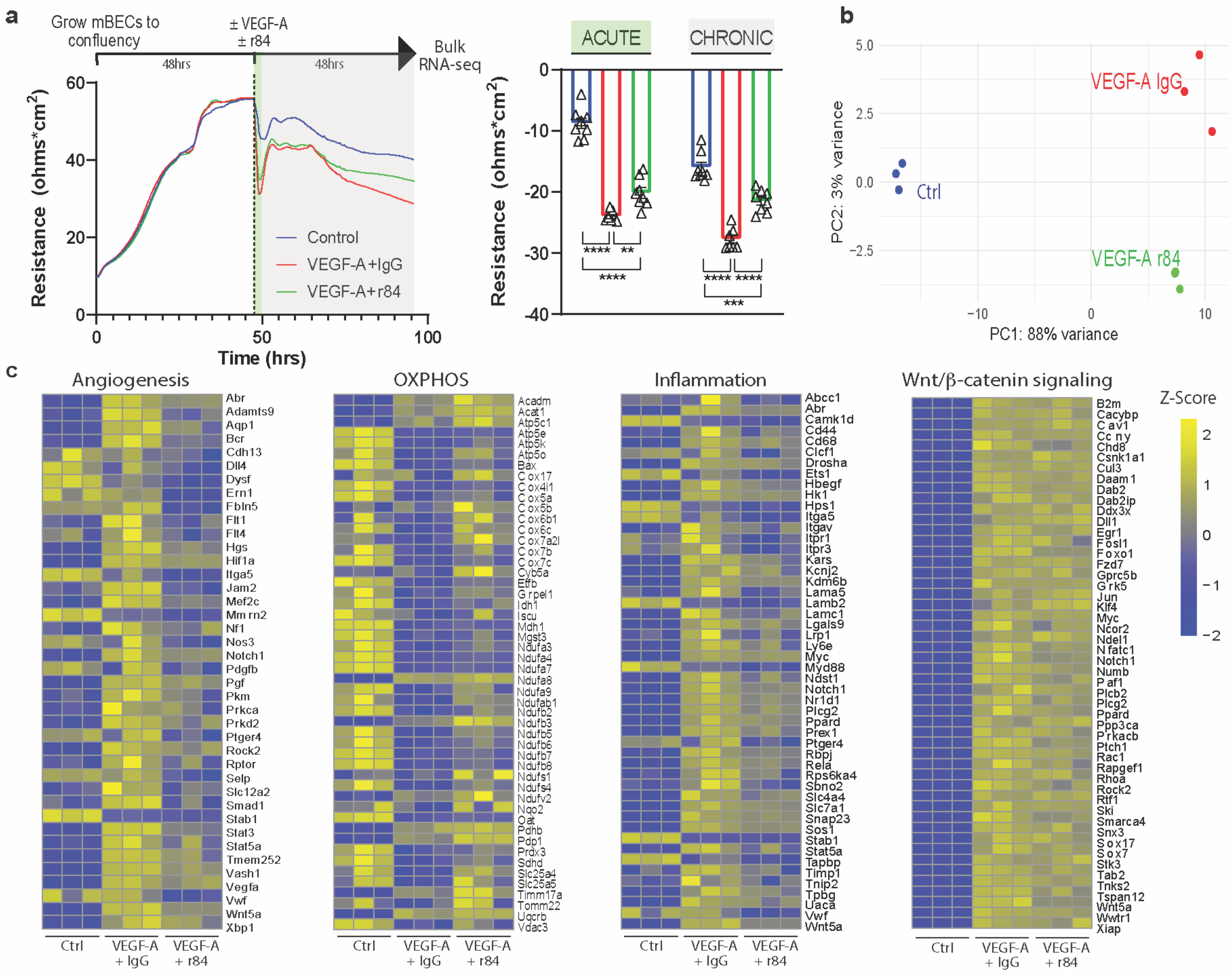


**Extended Data Figure 6.** ***In vitro*** **application of the humanized r84 antibody partially inhibits VEGF-A-mediated loss in barrier properties and transcriptional changes in primary mouse brain endothelial cells. a**) Trans-endothelial electrical resistance (TEER) over time of primary mouse brain endothelial cells (mBECs) following treatment with VEGF-A (100 ng/mL) in the presence of either human IgG control or r84 antibody at 100-fold molar excess. Time course is shown as average of 8 wells per condition. The experiment was repeated three times. The acute drop in TEER immediately following the treatment (green shading) and the chronic drop (gray shading) are quantified separately (*p<0.05, **p<0.01, ***p<0.001,****p<0.0001, unpaired two-tailed Student's t-test). **b**) Principal component analysis (PCA) of bulk RNA sequencing data collected 48 hours after treatment (n=3 biological samples per condition). **c**) Heat map visualization of differentially expressed genes (DEGs) driving pathway enrichment in VEGF-A + r84 versus VEGF-A + IgG-treated samples. Downregulated DEGs are related to "Angiogenesis"

(FDR q-val: 0.0012) and “Inflammation” (FDR q-val: 0.0011), whereas upregulated DEGs are related to “Oxidative Phosphorylation” (FDR q-val: 0.0033). There is no change in “Wnt/β-catenin signaling”. Control values are shown for reference.

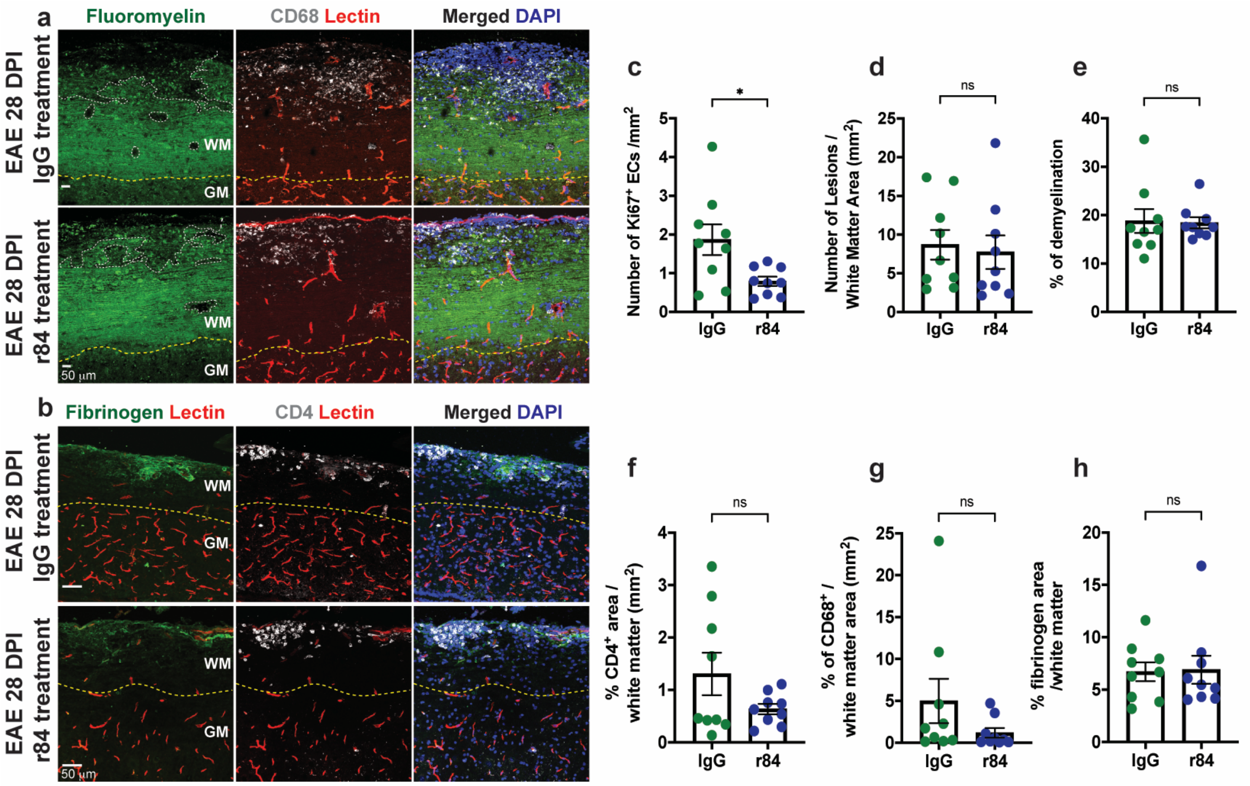


**Extended Data Figure 7. Treatment of $MOG_{35\text{-}55}$ EAE mice with the anti-VEGF-A monoclonal antibody r84 reduces endothelial cell proliferation, but does affect demyelination, immune cell infiltration and fibrinogen leakage in the spinal cord. a-b)** Sagittal sections of control IgG- and r84-treated (5 mg/kg) chronic (28 d.p.i.) EAE spinal cords were immunostained with antibodies against a) fluoromyelin and CD68 (marker of activated, phagocytic macrophages and microglia), and b) fibrinogen and CD4 (marker of T helper cells), as well as DAPI following cardiac perfusion with Lectin-A594 (marker of perfused vessels). **c)** Dotted bar graphs of the quantification of $Ki67^+$ ECs numbers / $mm^2$ reveal significantly lower EC proliferation in r84-treated compared to the IgG-treated control spinal cords (n = 9 mice/condition). **d-h)** Dotted bar graphs show no significant changes in the number of white matter lesions (d), area of demyelination (e), number of infiltrating $CD4^+$ T helper cells (f), number of activated microglia and infiltrating macrophages, and h) fibrinogen leakage between IgG-treated and r84-treated chronic EAE spinal cords.

### II. Extended Data Tables and Legends

**Extended Data Table 1**. Batch structure for single-cell RNA-seq experiments (sheet 1). Gene lists used for the GSEA analysis (sheet 2). Results of differential expression and GSEA analysis between acute vs. CFA/healthy aECs (sheet 3); acute vs. CFA/healthy cECs (sheet 4); chronic vs. CFA/healthy cECs (sheet 5); acute vs. CFA/healthy venule ECs (sheet 6); chronic vs. CFA/healthy venuleECs (sheet 7); acute vs. CFA/healthy vein ECs (sheet 8); chronic vs. CFA/healthy vein ECs (sheet 9); r84- versus IgG-treated venous (vein and venule) ECs (sheet 10). Negative log2fc values mean that the gene is lower in the disease state compared to CFA/healthy controls, and positive values mean that the gene is higher in the disease state compared to CFA/healthy controls.

**Extended Data Table 2.** Results of differential expression and GSEA analysis between VEGF-A and IgG-treated versus control mouse primary brain endothelial cells (mBECs; sheet 1), VEGF-A and r84-treated versus control mBECs (sheet 2), and VEGF-A and r84-treated vs VEGF-A and IgG-treated mBECs (sheet 3).

| Target | Company / Catalog info | Dilution |
|---|---|---|
| Caveolin-1 | Abcam / ab18199 | 1/1000 |
| Vcam-1 | eBioscience / 14-1061-85 | 1/100 |
| Ki67 | Invitrogen / 14-5698-80 | 1/200 |
| Ki67 (SP6) | Abcam / ab16667 | 1/250 (mouse); 1/100 (human) |
| Mfsd2a | Dr. David Silver, Duke-NUS | 1/500 |
| αSMA | Sigma / C6198-100UL | 1/1000 |
| CD31 | BD Pharmingen / 553370 | 1/250 |
| Emcn | Invitrogen / 14-5851-82 | 1/500 |
| Fibrinogen | Lifespan Biosciences / LS-C150799 | 1:2000 |
| CD4 | BD Pharmingen / 553727 | 1:50 |
| EphB4 | R&D / AF446 | 1:100 |

| | | |
|---|---|---|
| hEGFL7 | R&D / AF3638 | 1/40 |
| hMCAM | R&D / AF932 | 1/20 |
| hCD31 | DAKO / M0823 | 1/100 |
| hCD68 (SPM13) | Novus Biologicals / NBP2-32831 | 1/200 |
| **Other reagents** | | |
| FluoroMyelin | Invitrogen / F34652 | 1:300 |
| Tomato Lectin, DyLight 488 | Invitrogen / L32470 | N/A |

**Extended Data Table 3. Antibody information and dilutions used for immunofluorescence staining in spinal cord tissue.**

| BioBank Code | Patient ID | Age | Sex | Patient info | Type of MS plaques | FIXATION |
|---|---|---|---|---|---|---|
| 2734 | 1 | 52 | F | Multiple sclerosis | Chronic plaques | Formalin-fixed tissue |
| 3102 | 2 | 70 | F | Multiple sclerosis | Chronic and active plaques with mild perivascular cuffing | Formalin-fixed tissue |
| 3115 | 3 | 79 | F | Multiple sclerosis | Chronic plaques | Formalin-fixed tissue |
| 3134 | 4 | 56 | M | Multiple sclerosis | Chronic and active plaques | Formalin-fixed tissue |
| 3164 | 5 | 72 | M | Multiple sclerosis | Chronic plaques | Formalin-fixed tissue |
| 4817 | 6 | 38 | F | Multiple sclerosis | Chronic and active plaques | Formalin-fixed tissue |
| 5292 | 7 | 70 | F | Multiple sclerosis | Chronic plaques | Formalin-fixed tissue |
| 5139 | 8 | 81 | M | Multiple sclerosis | Chronic plaques | Formalin-fixed tissue |

| | | | | | | |
|---|---|---|---|---|---|---|
| 5095 | 9 | 60 | M | Multiple sclerosis | Chronic plaques | Formalin-fixed tissue |
| 5102 | 10 | 60 | F | Multiple sclerosis | Chronic and active plaques | Formalin-fixed tissue |
| 5123 | 11 | 79 | F | Multiple sclerosis | Chronic plaques | Formalin-fixed tissue |
| 5154 | 12 | 63 | M | Multiple sclerosis | Chronic plaques | Formalin-fixed tissue |
| 1516 | 13 | 66 | M | non-neurological control | Chronic and active plaques | Formalin-fixed tissue |
| 1525 | 14 | 61 | M | non-neurological control | n/a | Formalin-fixed tissue |
| 1533 | 15 | 79 | F | non-neurological control | n/a | Formalin-fixed tissue |
| 2358 | 16 | 78 | M | non-neurological control | n/a | Formalin-fixed tissue |
| 2657 | 17 | 78 | F | non-neurological control | n/a | Formalin-fixed tissue |
| 3119 | 18 | 77 | F | non-neurological control | n/a | Formalin-fixed tissue |
| 5293 | 19 | 41 | F | non-neurological control | n/a | Formalin-fixed tissue |
| 5072 | 20 | 83 | M | non-neurological control | n/a | Formalin-fixed tissue |
| 5190 | 21 | 68 | M | non-neurological control | n/a | Formalin-fixed tissue |
| 4320 | 22 | 87 | M | non-neurological control | n/a | Formalin-fixed tissue |
| 4660 | 23 | 73 | F | non-neurological control | n/a | Formalin-fixed tissue |
| 5265 | 24 | 80 | M | non-neurological control | n/a | Formalin-fixed tissue |

| | | | | | | |
|---|---|---|---|---|---|---|
| 5023 | 1b | 50 | F | Multiple sclerosis | Multiple plaques at various stages of activity. | Fresh Frozen |
| 5288 | 2b | 57 | F | Multiple sclerosis | Chronic plaques | Fresh Frozen |
| 5308 | 3b | 54 | F | Multiple sclerosis | Chronic plaques | Fresh Frozen |
| 5320 | 4b | 74 | F | Multiple sclerosis | Chronic plaques | Fresh Frozen |
| 4130 | 5b | 67 | F | non-neurological control | n/a | Fresh Frozen |
| 3795 | 6b | 72 | F | non-neurological control | n/a | Fresh Frozen |
| 3482 | 7b | 79 | F | non-neurological control | n/a | Fresh Frozen |
| 3348 | 8b | 76 | F | non-neurological control | n/a | Fresh Frozen |

**Extended Data Table 4. Demographic data for multiple sclerosis (MS) and non-neurological control cases used in this study.**

| **Target** | **Species** | **Dilution** |
|---|---|---|
| *Egfl7* | Mouse | 1/50 |
| *Apln* | Mouse | 1/50 |
| *Mcam* | Mouse | 1/50 |
| *Ecm1* | Mouse | 1/50 |
| *Serpine1* | Mouse | 1/50 |
| *SERPINE1* | Human | 1/50 |
| *ECM* | Human | 1/50 |
| *CLDN5* | Human | 1/50 |

**Extended Data Table 5. Antisense DIG-labeled RNA probe information and dilutions used for RNA *in situ* hybridization and FISH in this study.**

### SUPPLEMENTARY REFERENCES

1 Kaufmann, M. *et al.* Identification of early neurodegenerative pathways in progressive multiple sclerosis. *Nat Neurosci* **25**, 944-955, doi:10.1038/s41593-022-01097-3 (2022).
